# An integrated transcriptomics–functional genomics approach reveals a small RNA that modulates *Bacteroides thetaiotaomicron* sensitivity to tetracyclines

**DOI:** 10.1101/2023.02.16.528795

**Authors:** Daniel Ryan, Elise Bornet, Gianluca Prezza, Shuba Varshini Alampalli, Taís Franco de Carvalho, Hannah Felchle, Titus Ebbecke, Regan Hayward, Adam M. Deutschbauer, Lars Barquist, Alexander J. Westermann

## Abstract

Gene expression plasticity allows bacteria to adapt to diverse environments, tie their metabolism to available nutrients, and cope with stress. This is particularly relevant in a niche as dynamic and hostile as the human intestinal tract, yet transcriptional networks remain largely unknown in gut *Bacteroides* spp. Here, we map transcriptional units and profile their expression levels in *Bacteroides thetaiotaomicron* over a suite of 15 defined experimental conditions that are relevant *in vivo*, such as variation of temperature, pH, and oxygen tension, exposure to antibiotic stress, and growth on simple carbohydrates or on host mucin-derived glycans. Thereby, we infer stress- and carbon source-specific transcriptional regulons, including conditional expression of capsular polysaccharides and polysaccharide utilization loci, and expand the annotation of small regulatory RNAs (sRNAs) in this organism. Integrating this comprehensive expression atlas with transposon mutant fitness data, we identify conditionally important sRNAs. One example is MasB, whose inactivation led to increased bacterial tolerance of tetracyclines. Using MS2 affinity purification coupled with RNA sequencing, we predict targets of this sRNA and discuss their potential role in the context of the MasB-associated phenotype. Together, this transcriptomic compendium in combination with functional sRNA genomics—publicly available through a new iteration of the ‘Theta-Base’ web browser (www.helmholtz-hiri.de/en/datasets/bacteroides-v2)—constitutes a valuable resource for the microbiome and sRNA research communities alike.

## INTRODUCTION

The evolutionary success of bacteria is based on their ability to adapt to changing environmental conditions and is reflected in the plasticity of their transcriptome. When conditions are favorable, bacteria replicate quickly with generation times often less than 30 minutes. Consequently, when conditions change for the worse, gene expression must adapt rapidly. Part of this plasticity is brought about by transcriptional control mechanisms. Sigma (σ) factors are accessory components of the RNA polymerase complex that recognize defined promoter sequences, thus modulating the specificity of transcription initiation to functionally connected gene sets (Feklistov et al., 2014). Transcription initiation may be further fine-tuned by transcription factors that bind specific sequence motifs in the genome, thereby recruiting or repelling the RNA polymerase holoenzyme to or from their target genes (Balleza et al., 2009; Mejia-Almonte et al., 2020).

Complementing protein-mediated transcriptional control, environmental and pathogenic bacteria employ small regulatory RNAs (sRNAs) (Wagner and Romby, 2015). Via binding to partially complementary sequences within target mRNAs, these riboregulators can post-transcriptionally modulate gene expression to help bacteria adapt their metabolism to the available nutrients (Bobrovskyy and Vanderpool, 2013) and cope with noxious stimuli (Holmqvist and Wagner, 2017). For example, several sRNAs are involved in antibiotic stress responses in phylogenetically unrelated pathogens, proposing these molecules as relevant drug targets to treat infectious diseases (Dersch et al., 2017; Lalaouna et al., 2014; Mediati et al., 2021).

Tight control of gene expression is a prerequisite for bacteria to thrive in the dynamic and hostile niches of the mammalian intestinal tract. Bacteria of the Gram-negative *Bacteroides* genus are universal members of the gut microbiota of healthy human adults (Wexler and Goodman, 2017). These bacteria occupy a hub position in the distal colon, influencing both host physiology and incoming enteric pathogens (Bornet and Westermann, 2022), and serve as reservoirs of antibiotic resistance genes within the gastrointestinal tract (Whittle et al., 2002). Consequently, knowledge of the regulatory mechanisms underlying their gene expression can help devise microbiota-centric interventions to correct intestinal disorders.

Our current understanding of transcriptional control mechanisms in *Bacteroides* spp. mostly derives from studying their metabolic potential. Encoded on distinct clusters of neighboring genes, these bacteria harbor numerous polysaccharide utilization loci (PUL) (Terrapon et al., 2018), which allow them to feed on dietary fiber as well as on host glycans (Grondin et al., 2017). As individual PULs are tailored to the consumption of specific substrates, PUL expression has to be tightly controlled and tied to the nutritional fluctuations that are commonly associated with the gut environment. Indeed, PULs typically comprise regulatory systems—encoded within the same locus—that spur PUL transcription when the corresponding carbon source is sensed. Lacking the stereotypical primary σ factor of Proteobacteria, *Bacteroides* spp. harbor a different ‘house-keeping’ σ factor, σ^ABfr^ (Bayley et al., 2000; Vingadassalom et al., 2005), as well as numerous extra-cytoplasmic function σ/anti-σ factor pairs (Martens et al., 2008; Martens et al., 2009) that are often part of PULs specific to host-derived glycans (Martens et al., 2008). Alternatively, SusR-family transcription factors (D’Elia and Salyers, 1996) and hybrid two-component systems (Sonnenburg et al., 2006; Sonnenburg et al., 2010), which combine sugar-sensing and gene regulatory functions into a single polypeptide, may activate the transcription of their cognate PUL operon. On a higher hierarchical level, a conserved cyclic AMP receptor protein (CRP)-like global regulator, acts on multiple PUL systems in parallel and coordinates *Bacteroides* carbohydrate utilization with other cellular processes (Schwalm et al., 2016; Townsend et al., 2020).

In contrast to transcriptional regulation, little is known about the post-transcriptional control networks in *Bacteroides* spp. While individual members of this genus are known to encode hundreds of sRNAs (Cao et al., 2016; Ryan et al., 2020a), a primary bottleneck is that the vast majority do not yet have a known molecular function. The global assignment of *Bacteroides* sRNAs to metabolic and stress regulons has been hampered by the lack of comprehensive carbohydrate- and stress-specific transcriptomics datasets. What is more, induced expression on its own may not necessarily reflect an RNAs importance for bacterial fitness under that condition. Genome-wide phenotypic screens in *Bacteroides* spp. have so far been restricted to the analysis of mutations within coding genes (Goodman et al., 2009; Liu et al., 2021; Wu et al., 2015), yet knowledge of fitness-contributing noncoding genes could help prioritize sRNAs for functional studies in these health-relevant bacteria. To date, only few *Bacteroides* sRNAs have been partially characterized by serendipity. This includes the sRNAs GibS (Ryan et al., 2020a) and DonS (Cao et al., 2016), which both control the expression of metabolic target genes, and RteR that suppresses the transfer of an antibiotic resistance-conferring mobile genetic element (Waters and Salyers, 2012). However, inactivation of none of these sRNAs has been associated with a robust fitness phenotype of the resulting deletion mutant, obscuring their importance for *Bacteroides*’ physiology.

Here, we set out to dissect global gene expression signatures in *Bacteroides thetaiotaomicron* type strain VPI-5482 under a range of host niche-related stresses and during growth on defined carbon sources. To this end, we applied a hybrid transcriptomics approach combining transcription start site (TSS) mapping via differential RNA-seq (dRNA-seq) (Sharma and Vogel, 2014) and conventional RNA-seq-based expression profiling over 15 defined *in vivo*-relevant conditions. From the resulting transcriptomic compendium, we (re-)annotated transcript boundaries, inferred stress- and carbon source-specific gene expression patterns, and identified novel noncoding RNAs in this bacterium. In an integrative approach, we used gene expression and mutant fitness data to link individual sRNAs to specific cellular processes. To demonstrate the value of our combined transcriptomics and genomics data, we focus on the previously uncharacterized sRNA MasB (BTnc201). Our findings assign MasB to the regulon governed by the CRP-like transcription factor BT_4338 and suggest that this sRNA is a post-transcriptional regulator of a conserved tetratricopeptide protein, with phenotypic consequences when *B. thetaiotaomicron* is exposed to translation-blocking compounds. Taken together, this multi-modal study substantially extends ‘Theta-Base’ (www.helmholtz-hiri.de/en/datasets/bacteroides-v2)—our interactive *Bacteroides* transcriptome database—and proposes fitness-affecting sRNAs in these important gut commensals for functional characterization.

## RESULTS

### B. thetaiotaomicron transcriptomics across 15 experimental conditions

Previously, we published a transcriptome map of *B. thetaiotaomicron* (Ryan et al., 2020a), yet this was solely based on experiments in nutrient-rich laboratory medium that falls short in reflecting *in vivo*-relevant conditions, composed of diverse stresses and nutritional variation. To overcome these prior shortcomings, we compiled a suite of *in vitro* conditions that mimic specific aspects of this bacterium’s host niche (Table 1). The large intestine exerts selective pressure on colonizing bacteria in the form of fluctuating pH levels (pH 5.5–7.0), heterogeneous oxygen tension, and the presence of secreted antimicrobial peptides and bile salts (Nugent et al., 2001; Ryan et al., 2020b; Singhal and Shah, 2020; Yao et al., 2018). Consequently, our suite of stress conditions includes moderate acidic pH (pH 6), aerobic shaking, exposure to hydrogen peroxide (H_2_O_2_), the bile acid deoxycholate or a bile salt mixture, and to the antibiotic gentamicin to which *Bacteroides* spp. are naturally resistant, as well as elevated (42°C) or reduced (28°C) temperature (Supplementary Fig. S1a-f). Additionally, bacteria were grown in minimal medium supplemented with defined simple sugars (glucose, arabinose, xylose, maltose, *N*-acetyl-D-glucosamine [GlcNAc]) or with porcine mucin glycans (Supplementary Fig. S1e, f). Total RNA was extracted from the respective cultures and either pooled and analyzed via dRNA-seq for comprehensive TSS mapping (as described in (Kroger et al., 2013)), or sequenced separately via conventional RNA-seq to profile conditional gene expression (Fig. 1a). In all cases, library preparation was generic, resulting in the detection of both protein-coding and noncoding RNAs.

**Table 1:** Experimental conditions analyzed via RNA-seq. MM, minimal medium; TYG, tryptone yeast extract glucose; LEP, late exponential phase; MEP, mid-exponential phase; ESP, early stationary phase.

| Supplement | Concentration/<br>condition | Medium | Growth phase | OD<sub>600</sub> | Time [h] | Reference |
| --- | --- | --- | --- | --- | --- | --- |
| Glucose | 0.5% | MM | LEP | 1 | 17 |  |
| Arabinose | 0.5% | MM | LEP | 1 | 17 |  |
| Xylose | 0.5% | MM | LEP | 1 | 17 |  |
| Maltose | 0.5% | MM | LEP | 1 | 17 |  |
| GlcNAc | 0.5% | MM | LEP | 1 | 23 |  |
| Mucin | 0.5% | MM | ESP | 0.2 | 15.5 |  |
| Starvation | n. a. | MM | LEP | 1 | 2 post media change | (Schofield et al., 2018) |
| Cold (28°C) | 28°C | TYG | MEP | 2 | 2 post stress induction |  |
| Heat (42°C) | 42°C | TYG | MEP | 2 | 2 post stress induction |  |
| Acid (pH6) | pH 6 | TYG | MEP | 2 | 2 post stress induction |  |
| Aerobe | 220 rpm, O<sub>2</sub> | TYG | MEP | 2 | 2 post stress induction |  |
| H<sub>2</sub>O<sub>2</sub> | 480 µM | TYG | MEP | 2 | 2 post stress induction |  |
| Deoxycholate | 0.2 mg/mL | TYG | MEP | 2 | 2 post stress induction | (Liu et al., 2021) |
| Bile salts | 0.5 mg/mL | TYG | MEP | 2 | 2 post stress induction | (Liu et al., 2021) |
| Gentamicin | 200 µg/mL | TYG | MEP | 2 | 2 post stress induction |  |

**Figure 1:**
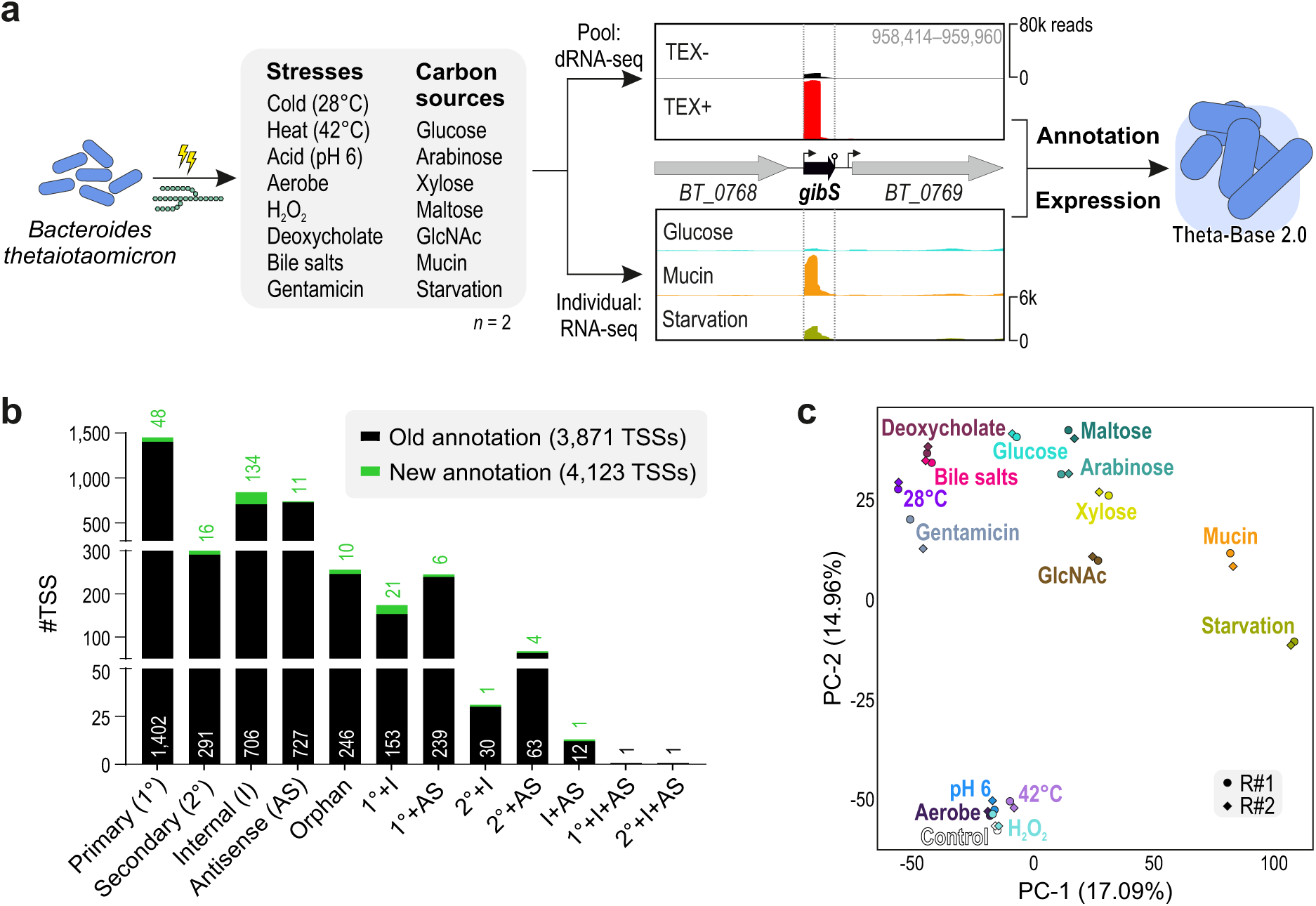
Comprehensive transcriptome annotation and gene expression profiling of *B. thetaiotaomicron*. a, Experimental outline. *B. thetaiotaomicron* type strain VPI-5482 was grown either in rich TYG medium to mid-exponential phase (OD_600_ 2.0) followed by a 2-hour exposure to the indicated environmental stresses or in minimal medium supplemented with the indicated carbon sources to late-exponential phase (OD_600_ 1.0) or—in case of mucin—early stationary phase (OD_600_ 0.2). RNA samples were either pooled for comprehensive TSS mapping via dRNA-seq or analyzed separately via conventional RNA-seq. TEX, Terminator exonuclease. **b,** Refined TSS annotations. See Methods for definitions of the different categories. **c,** Principle component analysis of all RNA-seq samples analyzed.

The pooled cDNA sample was sequenced to ∼40 million reads, i.e. twice the depth previously considered sufficient to annotate the transcriptome of *Salmonella enterica* serovar Typhimurium to saturation (Kroger et al., 2013). We analyzed the resulting data using the ANNOgesic pipeline (Yu et al., 2018) and collectively mapped the position of 4,123 TSSs across the *B. thetaiotaomicron* chromosome and plasmid (RefSeq IDs: NC_004663; NC_004703). Comparing these results to our previously mapped TSSs, when *B. thetaiotaomicron* grew in rich TYG medium in early or mid-exponential, and stationary phase (Ryan et al., 2020a), we found 252 extra unique TSS annotations contributed by the 15-condition pool (Fig. 1b, Supplementary Fig. S2a and Supplementary Table 6). Likewise, the number of transcription termination sites (TTSs)—predicted by a combination of read coverage drop and likelihood to fold into an intrinsic terminator hairpin (see Methods)—increased by 86 (Supplementary Fig. S2b). We also updated the annotation of operon structure predictions (Supplementary Fig. S2c). Of these 522 operons, two thirds are dicistrons. On the opposite side of the size spectrum, five operons encompass more than ten genes, with two of the largest encoding the capsular polysaccharide 4 (comprising of 13 genes) and a set of ribosomal factors (32 genes). Lastly, to better interpret *Bacteroides* transcriptomic features, we integrated our refined transcript boundary annotations with a map of invertible DNA regions (‘invertons’) obtained by the application of the PhaseFinder algorithm (Jiang et al., 2019) to the *B. thetaiotaomicron* genome. Of the resulting 1,997 inverted repeats, 569 contain potential promoters (i.e., involve sequences within a 50 bp window upstream of a mapped TSS). The latter may represent sites contributing to *Bacteroides* phase-variable transcription initiation (Jiang et al., 2019) and are relevant to dissect gene expression heterogeneity in this organism.

### Bacteroides stress response signatures

The conventional RNA-seq libraries were sequenced to 10–15 million reads per sample, as per general guidelines for bacterial differential expression analysis (Haas et al., 2012). Of the 5,442 coding sequences in the *B. thetaiotaomicron* genome, 5,137 (94.4%) were expressed (>10 reads/sample) in at least one of the 15 experimental conditions. Biological replicate samples clustered closely (Fig. 1c), indicating the absence of major batch effects. To aid in interpretation of the transcriptome dataset, we compiled functional information by merging KEGG and GO-term annotations with manually collated gene sets and regulons retrieved from literature research (see Methods and Supplementary Table 8).

Pairwise comparisons of brief stress exposures to an unstressed control sample (Supplementary Fig. S5; Supplementary Table 7) and an ensuing gene set enrichment analysis (Supplementary Fig. S3a) revealed *Bacteroides* transcriptomic responses to environmental cues. Heat, mild acidic pH, and aerobic exposure triggered only few, yet specific expression changes (Fig. 2a; Supplementary Fig. S5b-d). Stress-specific “marker genes” inferred from the literature showed the anticipated alterations (Lu and Imlay, 2021; Mishra and Imlay, 2013), including the induction of *BT_1606* (encoding cytochrome C peroxidase) and *BT_1456* (thioredoxin) when cultures were shifted to aerobic conditions (Supplementary Fig. S5d). Brief exposure to hydrogen peroxide did not induce any significant changes in expression as compared to the unstressed culture, probably because the selected peroxide concentration (480 µM; Supplementary Fig. S1b) was relatively low. In contrast, substantial transcriptomic reprogramming was observed when bacteria faced cold, bile, or sub-lethal antibiotic stress (Fig. 2a, c; Supplementary Fig. S5a, f). When exposed to deoxycholate or a bile salt cocktail, the most highly expressed operons included *BT_2792*–*BT_2795* and *BT_0691*–*BT_0692* (Fig. 2c; Supplementary Fig. S5e, g) that are both known to contribute to *Bacteroides* fitness under bile stress (Bechon et al., 2022; Liu et al., 2021). In case of the *BT_2792*–*BT_2795* operon that encodes a bile salt tolerance-conveying efflux pump (Liu et al., 2021), we observed a TSS and alternative start codon 60 nt downstream of the annotated one (red and black ‘ATG’ in Fig. 2d). Since the previously annotated N-terminus was not supported by any sequencing reads (Fig. 2d, bottom), we re-annotated *BT_2795* accordingly. During the course of this work, a new annotation update of the VPI-5482 genome (sequence ID NC_004663.1) has been released by NCBI that supports this observation.

**Figure 2:**
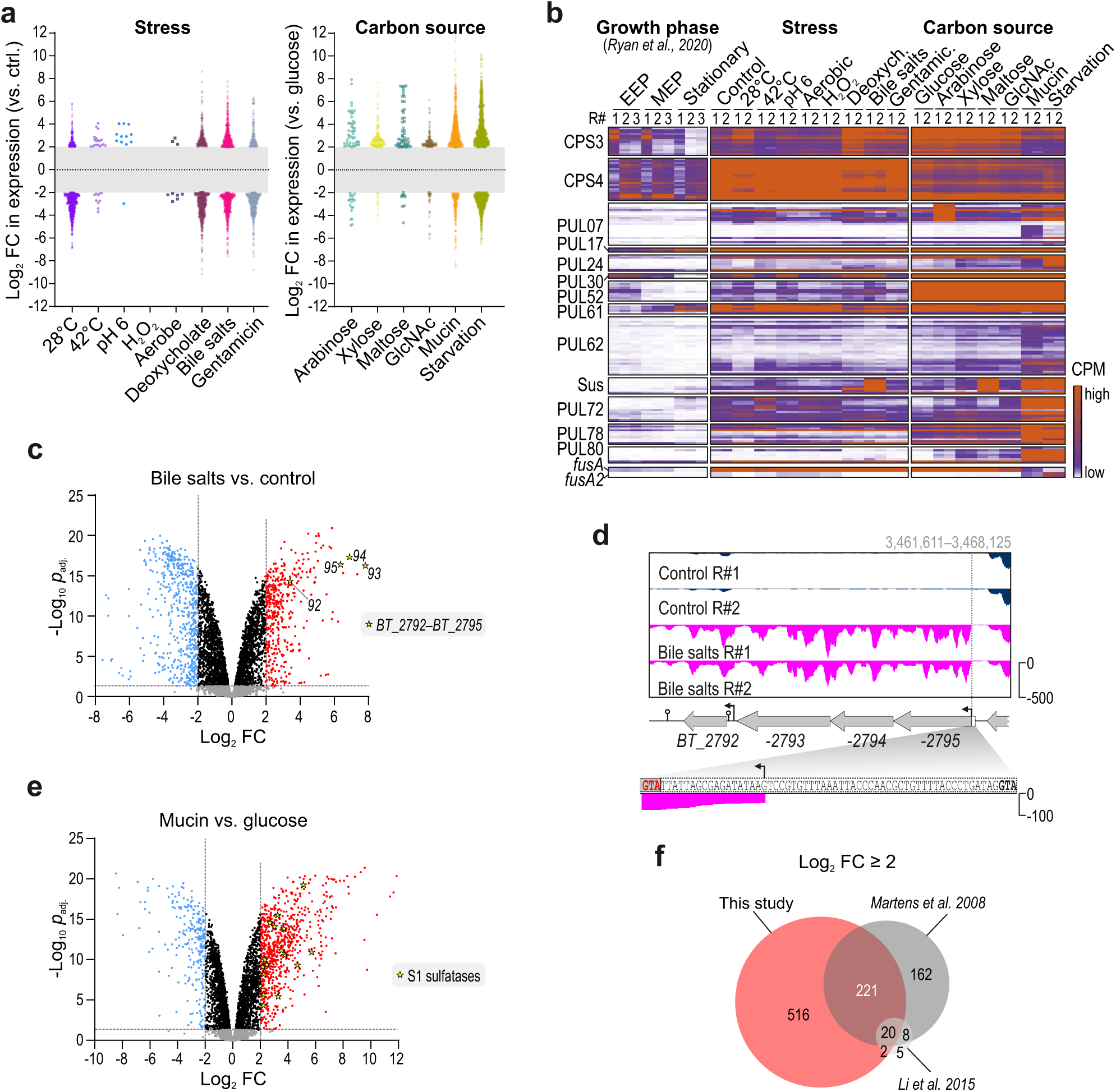
Stress response- and carbon source-specific expression of *B. thetaiotaomicron* genes. **a,** Bee swarm plots indicate global expression changes of *B. thetaiotaomicron* genes upon environmental shocks or growth in presence of the indicated sole carbon sources. Each dot denotes a bacterial gene that was up-(log_2_ FC >2, FDR <0.05) or downregulated (log_2_ FC <-2, FDR <0.05) as compared with the unstressed control condition in TYG medium, the growth in minimal medium plus glucose or, in case of starved bacteria, growth in rich TYG, respectively. Gray center bars delimit log_2_FC values that are not significant as per the above definition. **b,** Heat map showing the abundance of transcripts belonging to CPS3 or -4, or are derived from selected PULs or from elongation factor G-encoding mRNAs *BT_2729* (*fusA*) and *BT_2167* (*fusA2*). Growth phase-dependent expression data stem from (Ryan et al., 2020a). EEP, early exponential phase; MEP, mid-exponential phase; CPM, counts per million. The complete data for all eight CPSs and 96 PULs, as well as the CRP regulon are given in Supplementary Fig. S4. **c,** Volcano plot of differentially expressed genes when bacteria were exposed to bile salts compared with an unstressed control condition. Yellow asterisks denote genes of an operon coding for a bile salt tolerance-conveying efflux pump (Liu et al., 2021). **d,** Reannotation of the *BT_2792*–*BT_2795* operon was prompted by a newly discovered TSS downstream of its originally annotated start codon (bold black; proposed actual start codon in red). **e,** Volcano plot showing gene expression changes as *B. thetaiotaomicron* feeds on mucin compared with glucose. S1 sulfatase genes (https://doi.org/10.1101/2020.11.20.392076) are indicated by yellow asterisks. **f,** Venn diagram displays the overlap of mucin-specific gene induction between our dataset and those obtained in (Martens et al., 2008) and (Li et al., 2015).

*Bacteroides* genomes harbor multiple capsular polysaccharide (CPS) loci, allowing these bacteria to alter their surface structure. Different CPSs have varying immunogenicity and are composed of different outer membrane proteins that can be hijacked as adhesion receptors by bacteriophages. CPS switching therefore is a means for *Bacteroides* to evade host immunity and phage attack (reviewed in (Porter and Martens, 2017)). Invertible promoters—included in our inverton map (see above)—result in phase-variable expression of individual CPS loci (Porter et al., 2020). Of the eight CPSs of *B. thetaiotaomicron* VPI-5482, CPS3 and CPS4 were dominant during *in vitro* growth, recapitulating previous findings (Porter et al., 2017). Interestingly, however, the dominantly expressed type switched from CPS4 to CPS3 when bacteria were exposed to the secondary bile acid deoxycholate, and—although less prominently—when exposed to the bile salt mixture or gentamicin and when shifted to 28°C (Fig. 2b). As shown before in colonized mice (Porter et al., 2017), CPS3 expression increased transiently after a diet switch from low to high fiber, while being replaced by other CPSs (CPS4, -5, and -6) on the long term. Thus, the CPS3-encoded capsule might be favored when *Bacteroides* face major lifestyle transitions and respond to, for example, a transient influx of bile following digestion. Altogether, these data corroborate and extend former observations of stress-induced adaptations of *Bacteroides* gene expression.

### Carbon source-specific gene expression patterns

To dissect *B. thetaiotaomicron* metabolic programs, we calculated differential gene expression upon growth in the different carbon sources relative to cultures feeding on glucose (Supplementary Fig. S5; Supplementary Table 7) and again determined enriched gene sets (Supplementary Fig. S3b). The number of significantly (logFC <-2 or >+2 and FDR <0.05) differentially expressed genes tended to increase with the complexity of the carbon sources (Fig. 2a). Generally, PUL expression responded to known substrates, adding further confidence to our dataset (Fig. 2b). For example, arabinose specifically triggered PUL07 expression (previously found upregulated in presence of arabinan or pectic galactan (Martens et al., 2011)) and arabinose utilization operons (Schwalm et al., 2016) (Supplementary Fig. S5h). Maltose consumption was reflected by the specific upregulation of genes of the starch utilization system (Sus = PUL66; Supplementary Fig. S5j) (Cho et al., 2001; Reeves et al., 1997). In the presence of xylose, PUL61 genes and xylose utilization operons (Townsend et al., 2020) were induced (Supplementary Fig. S5i). PUL61 was also upregulated during growth on GlcNAc, as was PUL17.

When mucin was the sole carbon source, the high mannose mammalian *N*-glycan utilization system (PUL72; (Cuskin et al., 2015)) and host-derived mucin *O*-glycan-processing systems (PUL14, -78, and -80 (Briliute et al., 2019; Martens et al., 2008; Martens et al., 2011)) were strongly induced. PUL62, whose inducer is currently not known, was also upregulated under this condition, suggesting that this PUL responds to mucin-derived glycans. S1 sulfatase genes, required to process the heavily sulfated colonic glycoproteins (Luis et al., 2021), belonged to the most highly mucin-induced genes (Fig. 2e). Overall, a substantial fraction of the mucin-activated genes identified here overlapped with genes previously found to be upregulated in *B. thetaiotaomicron* growing *in vitro* on a glycan mixture prepared from the porcine gastric mucosa (Martens et al., 2008) or colonizing the outer mucus layer of C57BL/6 mice (Li et al., 2015) (Fig. 2f).

Finally, when bacteria were deprived of nutrients, they activated the stringent response to reallocate resources away from growth and favor persistence (Supplementary Fig. S3b; downregulation of gene sets related to translation, peptidoglycan biosynthesis, and cell cycle). Starved bacteria further induced expression of *araM* (*BT_0356*) and its associated arabinose-utilizing PUL07 (Supplementary Fig. S5l), which is all in line with previous reports (Schofield et al., 2018; Schwalm et al., 2016).

The transcriptional master regulator of the CRP family, BT_4338, is engaged in metabolic gene expression control in *B. thetaiotaomicron* (Schwalm et al., 2016; Townsend et al., 2020). Global expression patterns of known BT_4338 regulon members (Townsend et al., 2020) indicated major gene expression changes, particularly when bacteria consumed mucin or were starved (Supplementary Figs. S4 and S5l, m), in accordance with previous reports (Schwalm et al., 2016; Townsend et al., 2020). For instance, expression of *fusA2* (*BT_2167*), which is an established BT_4338 target gene (Townsend et al., 2020) and encodes the alternative translation elongation factor G2 (EF-G2), was induced in mucin and peaked during carbon deprivation (Fig. 2b). The inverse expression pattern was observed for *fusA* (*BT_2729*) (Fig. 2b), encoding the canonical EF-G and not belonging to the BT_4338 regulon (Townsend et al., 2020). This corroborates previous reports (Han et al., 2022; Schwalm et al., 2016; Townsend et al., 2020) and further supports that *B. thetaiotaomicron* utilizes a distinct protein synthesis machinery during colonization of a sugar-deprived host niche.

In summary, the combined data corroborate former reports, but also extend our collective knowledge of conditional gene expression in *B. thetaiotaomicron*. For example, we predict a new mucin-specific PUL system (PUL62) and a switch in CPS expression under certain stress conditions. The transcriptomic information on PULs, CPSs, and any other genetic feature, inverton predictions, as well as our curated functional gene set annotations can be freely accessed and interrogated through the new iteration of Theta-Base at www.helmholtz-hiri.de/en/datasets/bacteroides-v2. Interested users can find the instructions for how to exploit the different features of the website in the Methods section.

### Conditional expression of noncoding genes

Unlike the situation for *B. thetaiotaomicron* protein-coding genes, for which various reference datasets exist, there is hardly any information in the literature with respect to stress- and metabolism-related expression of noncoding genes of this bacterium. Therefore, we next turned our focus to noncoding RNAs. Interestingly, and in contrast to the relatively small fraction of novel TSSs and TTSs gained from the pooled dRNA-seq experiment (Fig. 1b; Supplementary Fig. S2b), these new data resulted in a substantial increase in the number of *B. thetaiotaomicron* noncoding RNA candidates (Fig. 3a). For example, upon manual curation (see Methods) we confidently predict 44 new intergenic and 12 new 3’UTR-derived sRNAs, as well as 80 novel *cis*-encoded antisense RNA (asRNA) candidates. Using northern blotting, we validated seven randomly selected sRNAs and two asRNAs (Supplementary Fig. S6a). Generally, noncoding RNAs were strongly differentially expressed across our experimental dataset (Supplementary Fig. S6).

**Figure 3:**
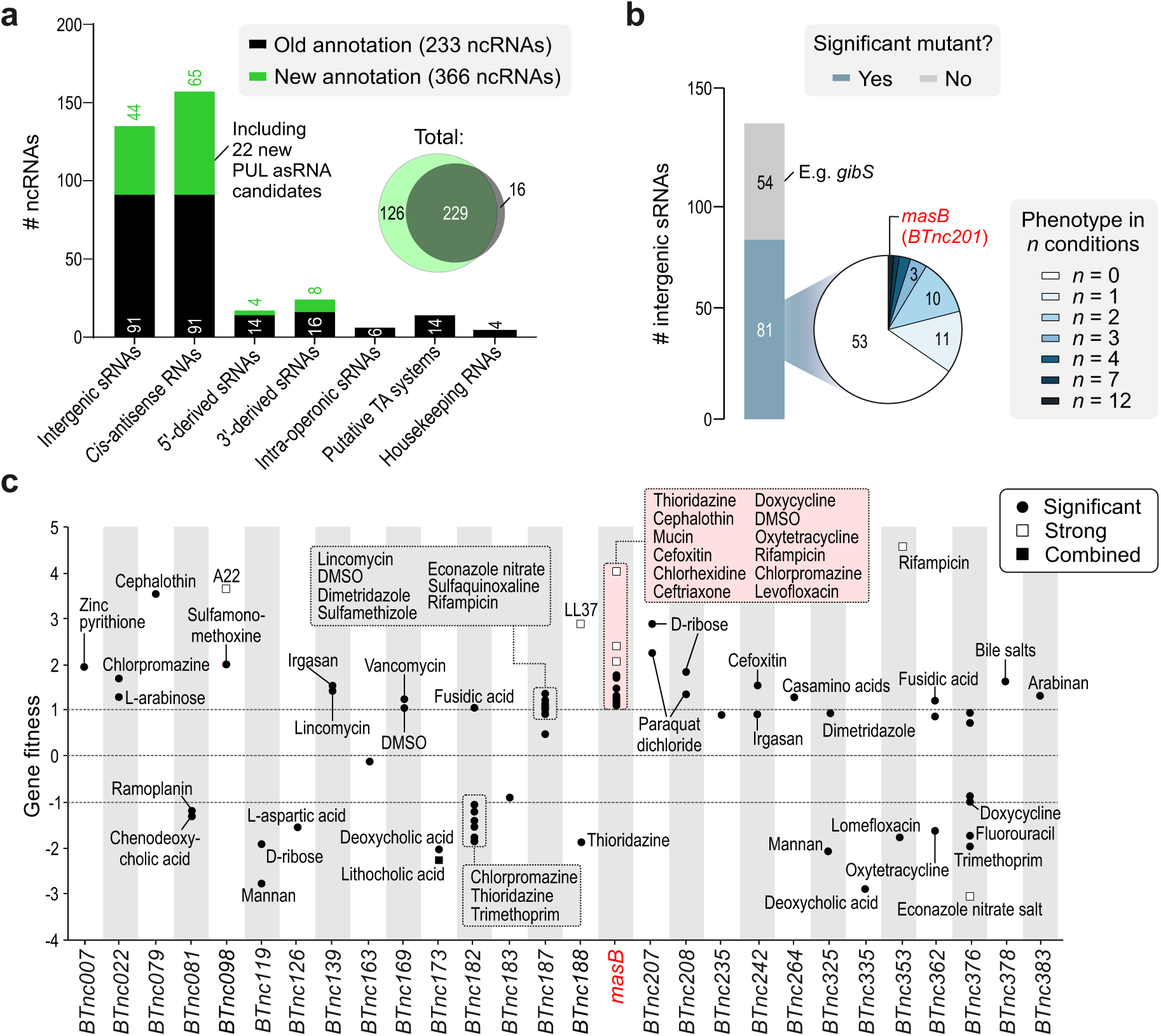
*Bacteroides* fitness-influencing sRNAs. **a,** Identification of novel noncoding RNA (ncRNA) candidates in *B. thetaiotaomicron*. For each class, the number of previously annotated ((Ryan et al., 2020a); over growth in rich medium; black) and the here-identified novel candidates (green) are plotted as a bar chart and the Venn diagram displays their overlap. **b, c,** Fitness data of *B. thetaiotaomicron* sRNA mutants, obtained by reanalyzing the dataset from (Liu et al., 2021). The sRNAs are grouped based on the number of significant fitness changes associated with their transposon disruption (**b**) and for sRNA mutants with at least one phenotype, the respective condition(s) and fitness score(s) are plotted (**c**). The categories in panel **c** (i.e., combined, significant, strong) are defined in the Methods section.

A previous study discovered a family of asRNAs divergently encoded to PUL operons, preferentially PULs with specificity to host-derived glycans (Cao et al., 2016). The observed anti-correlation in expression between several of these antisense/sense transcript pairs suggested the asRNAs to repress their cognate PUL—a mechanism validated exemplarily for one PUL-associated asRNA, termed DonS, in *Bacteroides fragilis* (Cao et al., 2016). In *B. thetaiotaomicron*, we recently annotated ten PUL-associated asRNAs and also noticed examples of asRNA/PUL anti-correlation over growth in rich medium (Ryan et al., 2020a). Our extended TSS mapping further increased the number of *B. thetaiotaomicron* PUL-associated asRNA candidates to 32 (Fig. 3a), and out of the ten new candidates, we exemplarily validated two by northern blotting (Supplementary Fig. S6a). What is more, the comprehensive metabolic expression data now allowed us to probe this anti-correlation phenomenon on a more global scale. We again found examples where an asRNA’s expression inversely mirrored the expression of its cognate PUL operon (Supplementary Fig. S6b, upper). Surprisingly however, we also observed counter-examples of positive correlation in individual asRNA/PUL pairs. This included a number of PUL-encoded asRNAs, which were maximally induced when bacteria grew on mucin or were starved; conditions wherein the divergently encoded PUL genes were also highly expressed (Supplementary Fig. S6b, lower). In other words, the extended transcriptomic data revealed a more nuanced picture of PUL-associated asRNAs than anticipated and further proposes this specialized class of noncoding RNAs for functional characterization.

### Phenotypes associated with B. thetaiotaomicron sRNA inactivation

To provide support for the involvement of individual noncoding RNAs in specific cellular processes, we reanalyzed an existing high-throughput transposon insertion sequencing (TIS) dataset from *B. thetaiotaomicron* grown under 490 defined *in vitro* conditions (Liu et al., 2021). For practical reasons, we here focused on standalone sRNA genes, encoded as independent transcriptional units (with their own cognate TSS and TTS) and not overlapping with other genetic features. Of these 135 intergenic sRNAs (including the 44 newly annotated ones; see above), 81 were represented in the transposon mutant library (Fig. 3b; Supplementary Fig. S7a). Mutants of 28 sRNAs (including seven newly annotated ones) exhibited a statistically significant fitness change (|t|>4) compared with the other mutants in the pool in at least one “successful” experiment (as defined in (Liu et al., 2021), i.e. an experiment in which a gene is represented by a sufficient number of barcode counts) (Fig. 3b, c; Supplementary Table 5). The majority of intergenic sRNAs affecting fitness showed a condition-specific phenotype when disrupted (Fig. 3b). However, in the case of a handful of sRNAs, disruption resulted in broader competitive fitness changes (i.e., mutants were affected under multiple conditions, with maximally 12 conditions for *masB*-deficient mutants).

In case of bacterial protein-coding loci, simultaneous measurement of mRNA levels and mutant fitness had previously revealed that the transcriptional responses to a specific environment are inaccurate predictors of gene deletion phenotypes (Jensen et al., 2017). We hypothesized that, in light of the same basic theme of gene expression control through antisense pairing, sRNA abundances might better correlate with bacterial fitness. Therefore, for the 95 intergenic sRNAs for which insertion mutants were obtained, we compared changes in expression (z-scores) with fitness alterations across the nine experimental conditions shared between the RNA-seq profiling and TIS datasets. However, based on this analysis relative expression of an sRNA is generally a poor predictor of its fitness contribution under a given condition, despite a few notable exceptions (sRNAs labeled in Supplementary Fig. S7b). For example, upregulation of certain sRNAs during aerobic shaking (BTnc139, BTnc227) coincided with the reduced fitness of the corresponding transposon mutants. Likewise, BTnc290 was activated in both arabinose and xylose, and its disruption led to growth defects when bacteria fed on these two carbon sources. Opposedly, several sRNAs that were highly induced in the presence of bile salts (including BTnc098, BTnc349, and BTnc378) counteracted bacterial resilience to this stress.

The combined expression and fitness data for *Bacteroides* sRNAs open leads for future functional investigation. To facilitate exploitation of these rich datasets, the fitness data for noncoding RNA mutants were also made publicly available for comparative analysis through Theta-Base. In the following, we followed up on the MasB sRNA, whose inactivation led to the highest number of significant fitness phenotypes amongst all intergenic sRNA mutants.

### MasB: an antibiotics susceptibility-conferring sRNA

MasB (previously BTnc201; renamed here for reasons to follow) is an ∼100 nt-long, narrowly conserved sRNA (Prezza et al., 2022). In *B. thetaiotaomicron*, the sRNA is encoded in between an operon composed of genes for a Na^+^/H^+^ antiporter (*BT_3638*), a thiazole biosynthesis adenylyltransferase (*thiF*), and a lipoprotein-releasing system ATP-binding protein (*BT_3640*), and a divergently encoded operon comprising a Na^+^-dependent transporter (*BT_3641*) and a hypothetical gene (*BT_3642*) (Fig. 4a). Relative to the transcripts from these flanking genes, MasB accumulated to very high steady-state levels under all of the experimental conditions tested here, but peaked when bacteria were starved (Fig. 4a, upper; Supplementary Fig. S6b).

**Figure 4:**
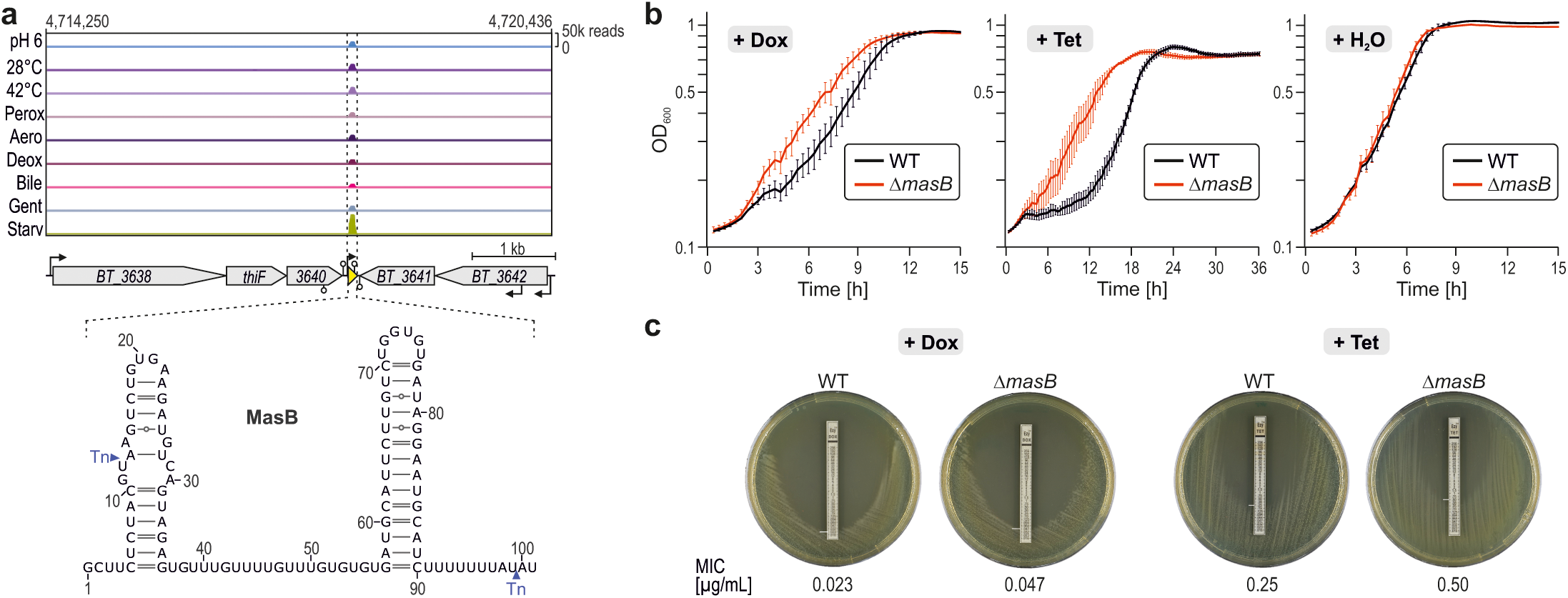
Genetic depletion of *masB* confers *B. thetaiotaomicron* enhanced tolerance of doxycycline and tetracycline. **a,** Upper: MasB expression in differently stressed *B. thetaiotaomicron* cultures. Shown are representative read coverages across the *masB* genomic locus and flanking genes. Lower: Experimentally determined secondary structure of MasB (Prezza et al., 2022). The positions of the inserted transposons are indicated by blue arrowheads. The insertion at position 12/13 results in a functional *masB* null mutant, while the insertion at position 99/100 is unlikely to affect this sRNA’s activity. **b,** Growth curves of a *B. thetaiotaomicron* Δ*masB* mutant (red) and its isogenic wild-type (black) in TYG, either supplemented with 0.016 µg/mL of doxycycline (left), with 0.05 µg/mL of tetracycline (middle), or water as the vehicle control (right). Plotted are the means +/-SD from three biological replicate experiments, each comprising technical duplicates. **c,** Doxycycline and tetracycline strip assays to determine the minimal inhibitory concentration (MIC), which is defined as the point on the antibiotic strip that is intersected by the growth inhibition ellipse. Representative images from three biological replicate experiments are depicted.

The TIS analysis suggested that MasB influences *B. thetaiotaomicron* fitness under various conditions, especially promoting growth upon exposure to diverse antibiotics and antimicrobials (Fig. 3c). This included enhanced fitness of *masB* disruption mutants during exposure to tetracycline derivatives (oxytetracycline, doxycycline hyclate), which target the bacterial ribosome. We confirmed this phenotype using a clean deletion mutant of this sRNA and increasing concentrations of doxycycline, both during growth in liquid medium (Fig. 4b; Supplementary Fig. S7c) and on solid agar (Fig. 4c). Similar effects were observed when exposing the strains to conventional tetracycline (Fig. 4c; Supplementary Fig. S7d). The fitness effects were stress-specific, as the mutant grew indistinguishably from an isogenic wild-type strain in vehicle-treated control cultures (Fig. 4b). Northern blotting excluded major changes in MasB steady-state levels when bacteria were exposed to either doxycycline or tetracycline (Supplementary Fig. S7e). Based on these results we concluded that MasB confers *Bacteroides* sensitivity to ribosome-targeting antibiotics of the tetracycline family. We hence name this sRNA MasB, for ‘<u>m</u>odulator of <u>a</u>ntibiotics <u>s</u>usceptibility in *<u>B</u>acteroides*’.

### MAPS identifies the mRNA for a tetratricopeptide domain protein as a MasB target

Our transcriptomics dataset lends itself for coexpression analysis to obtain insight into cellular regulatory circuits. Here, as an illustrative use case, we performed coexpression analysis for MasB (Fig. 5a). The result prompted us to explore the hypothesis that transcription of this sRNA might be governed by BT_4338—the CRP-like transcription factor controlling carbohydrate utilization in *B. thetaiotaomicon* (Schwalm et al., 2016; Townsend et al., 2020). Indeed, closer inspection of publicly available chromatin immunoprecipitation and sequencing (ChIP-seq) data revealed that one of the most significant BT_4338 binding sites in the *B. thetaiotaomicron* chromosome is located upstream of the *masB* gene (peak ID #598 in (Townsend et al., 2020)). This suggests that BT_4338 might act as a transcriptional activator of MasB, but further analysis is needed to confirm this assumption.

**Figure 5:**
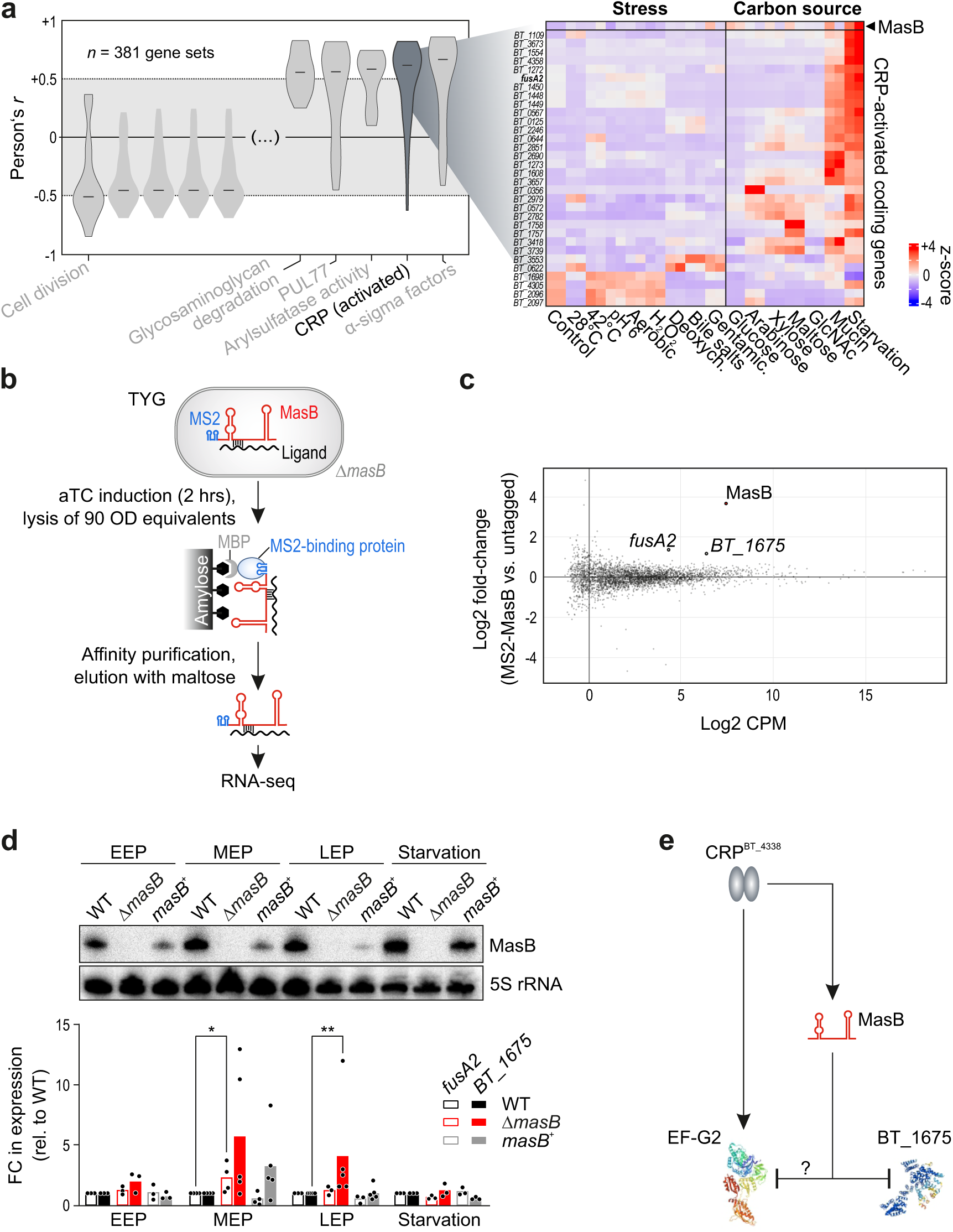
MasB is assigned to the CRP regulon and targets the mRNA of a conserved tetratricopeptide domain protein. **a,** MasB as part of the CRP regulon. Left: violin plots of Pearson’s correlation coefficients between MasB expression and that of annotated *Bacteroides* gene sets (comprised of ≥10 individual transcriptional units) sorted from left (lowest *r*) to right (highest *r*). Out of all 381 gene sets included in this analysis, the five most positively and negatively correlated ones (highest and lowest median *r*, respectively) are shown. Gene sets with |median *r*| > 0.5 are labeled with their name. The CRP-activated regulon (as defined in (Townsend et al., 2020)) ranked second. Right: heat map showing the expression of MasB and CRP-induced targets across the set of 15 different experimental conditions. **b,** Schematic of the MAPS workflow. *B. thetaiotaomicron* expressing an MS2-tagged version of MasB and an untagged control strain are grown to mid-exponential phase before induction of the sRNA’s expression using 200 ng/mL of aTC for 2 h. The resulting cell lysates are added to an amylose resin chromatography column to which a fusion of the maltose-binding protein (MBP) and the MS2 phage protein was attached. The MS2-tagged sRNA complex is captured by the MS2-MBP fusion on the column. The bound sRNA and its RNA ligands are subsequently eluted from the column through the addition of maltose and sequenced to identify binding partners of MasB. **c,** MD plot showing the log_2_ fold-change and average abundance of each gene in the MS2-MasB-expressing vs. untagged control strain. **d,** Upper: northern blot showing MasB abundance in early (EEP), mid-(MEP), late exponential phase (LEP), and upon starvation for the indicated strains. 5S rRNA was used as the loading control. Lower: qRT-PCR measurement of *fusA2* (outlined bars) and *BT_1675* (filled bars) mRNA abundances normalized to 16S rRNA in the same strains and growth conditions. Significance was calculated using a Mann-Whitney U test (*: *p* < 0.05; **: *p* < 0.01). **e,** Schematic of the working model of MasB-mediated regulations. The CRP-like transcription factor BT_4338 activates MasB transcription, which in turn post-transcriptionally represses *BT_1675* expression. Besides, the sRNA may also affect the mRNA for the elongation factor G2, whose transcription is also governed by BT_4338. Depicted is the cryo-electron microscopy structure of *B. thetaiotaomicron* EF-G2 (10.2210/pdb8DMF/pdb) and the AlphaFold (Jumper et al., 2021) representation of BT_1675 (UniProt ID Q8A750). The question mark indicates that the MAPS-predicted targeting of *fusA2* mRNA by MasB has not been validated.

To search for MasB targets, we here established MS2 affinity purification and sequencing (MAPS) technology (Lalaouna et al., 2017) in *B. thetaiotaomicron* (Fig. 5b). We fused a tandem MS2 aptamer to the 5’ end of MasB, as *in silico* RNA folding suggested this would maintain the native sRNA secondary structure (Supplementary Fig. S8a). The strain expressing the tagged version of the sRNA from an inducible promoter in the Δ*masB* background did not show a growth defect as compared with wild-type bacteria (Supplementary Fig. S8b, c). A strain expressing the untagged sRNA was included as a control (Supplementary Fig. S8d). After optimizing the affinity purification protocol for *Bacteroides* (see Methods; Supplementary Fig. S8e), we subjected the copurified RNA samples from two independent pulldown experiments to cDNA library preparation and sequencing (Supplementary Fig. S8f). The top-enriched transcripts in the MS2-MasB copurifications relative to the untagged background control were *BT_1675*, encoding a conserved tetratricopeptide domain protein, and the *fusA2* mRNA (*BT_2167*) that encodes the alternative ribosomal factor EF-G2 (Fig. 5c; Supplementary Fig. S9a).

Despite the relatively harsh purification conditions of our MAPS approach, leading to partial degradation of less protected RNA stretches (Supplementary Fig. S8e), we did not observe single sharp peaks in read coverage across the copurified transcripts that would pinpoint MasB binding sites (Supplementary Fig. S9a). To narrow in on potential targeting regions, we thus applied *in-silico* prediction of partially complementary sequences between the sRNA and its presumed targets using the IntaRNA algorithm (Busch et al., 2008; Wright et al., 2014). For *fusA2* no convincing binding site was found, however, *in-silico* prediction suggested that the *BT_1675* mRNA might be targeted ∼40 nt downstream of its translation start codon (Supplementary Fig. S9b). Confidence for this predicted intermolecular base-pairing came from the fact it involves a putative ‘seed’ region of MasB, which largely falls within the weakly structured central portion of this sRNA (Fig. 4a, lower). Indeed, electrophoretic mobility shift assays (EMSAs) confirmed MasB binding to the 5’ region of *BT_1675* mRNA *in vitro* (Supplementary Fig. S9c).

To test for an effect of MasB on the steady-state levels of *BT_1675* mRNA, we grew *B. thetaiotaomicron* wild-type, Δ*masB*, and *masB*^+^ (a *trans*-complementation strain of MasB) cultures in TYG or starved them for 2 h in carbon source-deprived minimal medium. We then extracted total RNA and subjected the samples to northern blot and qRT-PCR analyses (Fig. 5d). The *BT_1675* mRNA level was ∼5-fold de-repressed in the absence of MasB, yet exclusively so in mid- and late-exponentially growing bacteria. This suggests the exponential phase as the critical time window when MasB exerts a negative effect on *BT_1675* and reflects the observed growth phenotype of the sRNA deletion mutant (Fig. 4b). *Trans*-complementation of MasB reverted *BT_1675* expression to near wild-type levels (Fig. 5d). Similar observations—yet at lower scale— were made for the mRNA levels of *fusA2* (Fig. 5d). Based on these data, we suggest MasB to act as a riboregulator of the conserved tetratricopeptide domain protein BT_1675 through direct binding to the 5’ region of the corresponding mRNA.

## DISCUSSION

*Bacteroides* spp. are predominant members of the healthy human gut microbiota and must adapt to a variety of different nutrients and stressors (Wexler and Goodman, 2017; Wexler, 2007). Transcriptomics has proven invaluable for our understanding of the molecular basis of *Bacteroides*’ activities in this complex and dynamic niche. For example, RNA-seq in colonized mice revealed the transcriptional reprogramming of *Bacteroides sartorii* in face of host immune responses, spatial gene expression patterns of *B. fragilis* colonizing different mucosal and luminal gut niches, and the expression signature of *B. thetaiotaomicron* attached to the outer mucus layer (Becattini et al., 2021; Donaldson et al., 2020; Li et al., 2015). Disentangling *in-vivo* gene expression patterns gets complicated by the multitude of overlapping stimuli that the microbes are simultaneously exposed to in their natural niche. However, pinpointing the precise triggering factors that induce the expression of certain gene sets is important to understand the underlying regulatory networks and to obtain a molecular handle to rationally interfere with these processes for the benefit of the human host.

To decompose host-adapted bacterial gene expression, specific responses to certain environmental cues can be reconstituted *in vitro*. For example, comprehensive transcriptome compendia exist for various bacterial pathogens grown in culture flasks, under defined infection-mimicking cues, such as ‘SalCom’ (Kroger et al., 2013), ‘AcenitoCom’ (Kröger et al., 2018), ‘PneumoExpress’ (Aprianto et al., 2018), and ‘PATHOgenex’ (Avican et al., 2021). Similarly, metabolic networks of aerobic model bacteria have been reconstructed from transcriptomic data (Brandes et al., 2012; Caglar et al., 2017; Kim et al., 2018). Despite anaerobic gut commensals also being exposed to changing environments—including variations in the levels of antimicrobials, oxygen tension, and dietary intake—similar conditional transcriptomic datasets have not previously been compiled for many medically relevant intestinal microbiota members, including *Bacteroides* spp.

Here, we comprehensively profiled transcriptomic plasticity of the emerging microbiota model species *B. thetaiotaomicron*. By sampling a suite of 15 defined metabolic and stress conditions, we substantially expanded the transcriptome annotation and conditional gene expression information of the type-strain VPI-5482. We find that certain stresses such as low temperature, bile stress, and gentamicin exposure induced a proportionally greater response than the apparently more tolerable heat, mild acidity, and oxidative stress. This argues that the latter conditions are less detrimental for *Bacteroides* and hence, require only specific adaptations in gene expression to cope with. For instance, thanks to the activities of cytochrome C peroxidase and thioredoxin, *B. thetaiotaomicron* can tolerate short exposure to atmospheric oxygen (Lu and Imlay, 2021; Mishra and Imlay, 2013).

Metabolic gene expression profiling revealed that the extent of transcriptional reprogramming increased with complexity of the carbohydrate source. Of note, the majority of differentially regulated genes during growth on GlcNAc, a key component of the core mucin structure, was similarly affected during growth on mucin, suggesting mucin-specific expression changes to be mostly due to the availability of GlcNAc. We further observed substantial overlap between the *Bacteroides* transcriptomic response to mucins and to nutrient deprivation, including the upregulation of several PULs and S1 sulfatases that are required to breakdown complex mammalian glycans. To put this into context, the lack of dietary fiber causes *B. thetaiotomicron* to switch its metabolism to the consumption of host-derived glycans, the most abundant of which are mucin glycans derived from the intestinal epithelium (Bjursell et al., 2006; Sonnenburg et al., 2005).

All these transcriptomic data can be explored through our open-access online resource ‘Theta-Base’ (www.helmholtz-hiri.de/en/datasets/bacteroides-v2). For example, when interested in a particular gene of unknown function, this resource can provide useful guidance as to the biological process the corresponding gene product is involved. That is, gene coexpression analysis can inform on the transcriptional activator of a gene and assign it to a larger regulon. Besides, also for routine molecular biology tasks such as cloning endeavors and probe-based transcript detection and quantification, knowledge of the precise transcript boundaries, promoter regions, and experimental conditions when a given RNA is present in the cell, can be exploited for the optimal design of experimental protocols. Lastly, our manually compiled and curated pathway annotation is expected to facilitate future functional studies in *B. thetaiotaomicron* and further *Bacteroides* species.

Importantly, we complemented the transcriptomic data with functional sRNA genomics, revealing specific phenotypes for 28 *B. thetaiotaomicron* sRNAs. These mutant fitness data, too, are accessible through the new iteration of Theta-Base, which thus features the first genome-wide phenotypic dataset of *Bacteroides* noncoding RNAs. The predictive power of this integrative transcriptomics–functional genomics approach is exemplified by our findings on the previously uncharacterized sRNA MasB. Coexpression analysis suggested this sRNA to belong to the BT_4338 (CRP) regulon, in line with previous ChIP-seq data (Townsend et al., 2020), and TIS predicted a fitness benefit of *masB* loss-of-function mutants during antibiotic stress—a phenotype that we validated for a clean *masB* deletion mutant exposed to low concentrations of tetracycline-class antibiotics.

At the molecular level, MasB targets the mRNA for the hypothetical tetratricopeptide repeat protein BT_1675, whose function is currently unknown. However, the same coexpression analysis approach we used for MasB (Fig. 5a) revealed a strong positive correlation (Pearson’s *r*: 0.91) between *BT_1675* and the GO term “unfolded protein binding”, comprising protein chaperones (Supplementary Fig. S10). Indeed, tetratricopeptide repeats are often contained in bacterial chaperones (Cerveny et al., 2013), suggesting that BT_1675 could belong to this category of proteins. The precise mode of action employed by MasB to regulate *BT_1675* expression and whether this regulation is responsible for—or contributes to—the observed antibiotic phenotype warrants further investigation.

Tetracycline-like antibiotics prevent the attachment of aminoacyl-tRNA to the ribosomal acceptor site, thereby inhibiting bacterial proliferation. A growing number of bacterial species are acquiring resistance to the activity of tetracyclines and one of the most widespread mechanisms of tetracycline resistance is associated with translation elongation factor G (EF-G)-like proteins that confer ribosomal protection (Chopra and Roberts, 2001; Connell et al., 2003; Li et al., 2013b; Spahn et al., 2001). In fact, *B. thetaiotaomicron* EF-G2 bears similarity (∼23% amino acid sequence identity) to one such protein, TetQ (Nikolich et al., 1992). It is thus tempting to speculate that the fitness advantage of the Δ*masB* mutant under tetracycline stress might at least partially be explained by MasB-mediated regulation of *fusA2* expression. Despite MAPS predicting *fusA2* mRNA as a MasB target candidate, our attempts failed to uncover an sRNA-mRNA interaction site and we only observed minor MasB-dependent effects on *fusA2* expression levels. Future analysis is therefore required to functionally dissect the *masB* phenotypes and to map the complete target suite of this sRNA.

Despite these current knowledge gaps, our present data highlight the relevance of MasB for antibiotic sensitivity in major human gut commensals. More generally, our study emphasizes the power of combining bacterial expression atlases with additional data modalities. Building on a brand-new visualization tool, Theta-Base 2.0 allows easy and intuitive interaction with our diverse datasets and constitutes a much-needed resource for the microbiome research community.

## EXPERIMENTAL PROCEDURES

### Bacterial strains and cultivation

A detailed list of strains, plasmids, and oligonucleotides used in this study can be found in Supplementary Table 1. *Bacteroides* strains were routinely cultured in an anaerobic chamber (Coy Laboratory Products) with an anaerobic gas mix (85% N_2_, 10% CO_2_, 5% H_2_), at 37°C. Routine cultivation of all strains was performed in TYG medium and on supplemented BHI plates. For a detailed description of media composition and culture conditions for the RNA-seq analysis, refer to Table 1 and Supplementary Table 1. Growth assays in the presence of diverse carbon sources were carried out in minimal medium (MM), supplemented with 0.5% of a suitable carbon source as follows. A single colony of wild-type *B. thetaiotaomicron* VPI-5482 (AWS-001) was inoculated into 5 mL of MM-glucose and incubated anaerobically for 24 h. One milliliter of this culture was centrifuged (4,500 rpm, 3 min) to pellet bacterial cells that were re-suspended in an equal volume of MM (without carbon source). This was subsequently used to inoculate (1:100 dilution) MM containing an appropriate carbon source and incubated for the indicated time, following which aliquots (∼4 OD each) were collected for RNA extraction. Stress response assays were performed in TYG medium as indicated below. A single colony of AWS-001 was inoculated into 5 mL of TYG medium and incubated anaerobically, overnight. It was then sub-cultured into 100 mL of TYG medium and grown to mid-exponential phase (∼7 h, OD = 2.0). This culture was sub-divided into 5 mL fractions corresponding to each stress condition and centrifuged to pellet bacterial cells as before. The pellet was then re-suspended in an equal volume of TYG containing the indicated concentration of a stressor and incubated for a further 2 h, following which samples were collected for RNA extraction.

### Total RNA purification and removal of genomic DNA

All samples were collected as biological duplicates. Total RNA was isolated by the hot phenol method as follows. Briefly, bacterial cultures containing a total of ∼4 OD equivalents of cells were collected and one-fifth volume of stop mix was added (5% vol vol^-1 water saturated phenol, pH >7.0, 95% vol vol^-1 ethanol) (Eriksson et al., 2003). Cell lysis was achieved by incubation with lysozyme (600 µL of 0.5 mg mL^-1) and SDS (60 µL of 10% solution) for 2 min at 64°C with the subsequent addition of NaOAc (66 µL of 3 M solution). Extraction with 750 μL phenol (Roti-Aqua phenol) was carried out at 64°C for 6 min followed by the addition of 750 μL of chloroform. Precipitation of RNA from the aqueous phase was achieved with twice the volume of ethanol and 3 M NaOAc (30:1) mix and incubated at –80°C overnight. The samples were then centrifuged and the pellets washed with ethanol (75% vol vol^-1), followed by resuspension in 50 µL of RNase-free water. Traces of genomic DNA were removed by treating ∼40 µg of total RNA with 5 U of DNase I (Fermentas) and 0.5 µL of Superase-In RNase Inhibitor (Ambion) in a reaction volume of 50 µL. Samples for dRNA-seq were prepared by pooling equimolar amounts (each 100 ng) of total RNA from each condition.

### cDNA library preparation and sequencing

For dRNA-seq, prior to synthesizing cDNA, pooled total RNA was fragmented with ultrasound (4 pulses of each 30 sec at 4°C) and then treated with T4 Polynucleotide kinase (NEB). The RNA of each sample was then divided in half, one of which was treated with terminator exonuclease (TEX) to enrich for primary transcripts. The RNA was then poly(A)-tailed using poly(A) polymerase and the 5’-PPP was removed using 5’ polyphosphatase (Epicentre). RNA adaptors were then ligated and synthesis of first strand cDNA was performed using M-MLV reverse transcriptase and oligo(dT) primers. The cDNA was subsequently amplified to a concentration of ∼10-20 ng µL ^-1, purified (Agencourt AMPure XP kit, Beckman Coulter Genomics) and fractionated in a size range of 200-600 bp. The libraries were deep-sequenced on an Illumina NextSeq 500 system using 75 bp read-length at Vertis Biotechnologie AG, Germany.

For conventional RNA-seq, samples were first rRNA-depleted using the Pan-Prokaryote riboPOOLs kit (siTOOLs Biotech). This involved incubation of 1 µg of total RNA with 100 pmol of rRNA-specific biotinylated DNA probes at 68°C for 10 min, followed by a shift to 37°C for 30 min in 0.25 mM EDTA, 2.5 mM Tris-HCl pH 7.5, and 500 mM NaCl. Depletion of rRNA-DNA hybrids was achieved by two 15 min incubation periods with streptavidin-coated magnetic Dynabeads MyOne C1 (0.45 mg ThermoFisher Scientific) in 0.25 mM EDTA, 2.5 mM Tris-HCl pH 7.5, 1M NaCl at 37°C. The samples were then purified using the Zymo RNA Clean & Concentrator kit along with DNase treatment (Zymo Research). Libraries were prepared with the NEBNext Multiplex Small RNA Library Prep kit for Illumina (NEB) according to the manufacturer’s instructions and the following modifications. Samples were fragmented at 94°C for 2.75 min as per the NEBNext Magnesium RNA Fragmentation Module (NEB) with subsequent RNA purification using the Zymo RNA Clean & Concentrator kit. The fragmented RNA was then 3’-dephosphorylated, 5’-phosphorylated, and decapped with 10 U of T4 PNK +/− 40 nmol of ATP and 5 U of RppH (NEB). After each step, RNA was purified as mentioned above. The fragmented RNA was then ligated to adapters (3’ SR and 5’ SR, pre-diluted 1:3 in nuclease-free water) and the cDNA amplified for 14 cycles. These barcoded libraries were purified using MagSi-NGSPREP Plus beads (amsbio) at a 1.8:1 ratio of beads to sample volume. Libraries were checked for quantity and quality using a Qubit 3.0 Fluometer (ThermoFisher) and a 2100 Bioanalyzer with the high sensitivity DNA kit (Agilent). Pooled libraries were then sequenced on the NextSeq 500 platform (Illumina) at the Core Unit SysMed of the University of Würzburg.

### Read processing and mapping

Generated reads were quality-checked using FastQC (v0.11.8) and adapters were trimmed using Cutadapt (v1.16) with Python (v3.6.6), using the following parameters: -j 6 -a Illumina Read 1 adapter=AAGATCGGAAGAGCACACGTCTGAACTCCAGTCA -a Poly A=AAAAAAAAAAA -- output=out1.fq.gz --error-rate=0.1 --times=1 --overlap=3 --minimum-length=20 --nextseq-trim=20 3_1. For the both sequencing data types (dRNA-seq and conventional RNA-seq), READemption (v0.4.5) was used to map reads to the *B. thetaiotaomicron* VPI-5482 reference genome (NC_004663.1) and plasmid (NC_004703.1). Details of the alignment statistics can be found in Supplementary Table 4.

### Transcriptome annotation

We used the ANNOgesic pipeline (v0.7.33) to update the annotations of TSSs, terminators, operons, and ncRNAs as previously described (Ryan et al., 2020a). In short, TSSs were identified by using the “TSSpredator” function that compares the relative enrichment of reads between TEX-treated and untreated libraries in order to identify enriched peaks that are characteristic of the protected 5’ ends of primary transcripts. TSSs were classified on the basis of this enrichment and distance, relative to a coding gene. Primary TSSs were identified as having the highest enrichment within a 300 bp region upstream of an ORF, while all other TSSs within this region were classified as secondary TSSs. Internal TSSs were defined as originating on the sense strand within a coding sequence, while antisense TSSs were those that originated on the antisense strand and overlapped with or within 100 bp flanks of a sense gene. All remaining TSSs were classified as orphan TSSs. In order to predict terminators, ANNOgesic utilizes two heuristic algorithms, one of which scans the genome for Rho-independent terminators (TransTermHP) and the other predicts terminators by detecting a decrease in coverage between two adjacent genes. Leveraging the wealth of these diverse transcriptomic datasets, we additionally utilized the operon detection function of ANNOgesic (default settings) to predict both operons and sub-operons based on the other detected features namely, TSSs, transcripts, and genes supplied as GFF (general feature format) files.

The prediction of noncoding RNAs was done using the “srna” function of ANNOgesic that first compares predicted transcripts to known RNAs within the sRNA database (all sequences were downloaded from BSRD (Li et al., 2013a)) and the non-redundant (nr) protein database (ftp://ftp.ncbi.nih.gov/blast/db/FASTA/). Candidates that were detected in the sRNA database were identified as sRNAs, while those contained in the nr-database were excluded from further analysis. All remaining transcripts were classified as intergenic sRNAs if they possessed a defined TSS, stable secondary structure (RNAfold normalized folding energy < -0.05), length within 30 to 500 nt, and did not overlap with any other genetic feature in either sense or antisense orientation. The prediction of *cis*-antisense RNAs was based on the same criteria along with the presence of an annotated gene on the opposing strand. UTR-derived sRNAs were classified as 5’, if they shared the TSS with an mRNA and were associated with a coverage drop or processing site before the CDS. Similarly, 3’ UTR-derived sRNAs were predicted by either a TSS or processing site in the 3’ UTR and a processing site or shared terminator with the mRNA. Intra-operonic sRNAs were associated with a TSS or processing site at the 5’ end and a coverage drop or processing site at the 3’ end. All sRNAs can be interrogated using either their ID (‘BTncxxx’) or—when applicable— their trivial name (for example ‘MasB’).

As with all predictions, manual validation is necessary to ensure accuracy and reproducibility of the data. Some of the important criteria for exclusion of a predicted noncoding RNA candidate were as follows: 1) no significant change in coverage between the TEX-treated and-untreated libraries; 2) part of a longer transcript without clear boundaries delineating the sRNA; 3) extensive overlap with an annotated terminator sequence; 4) sequence overlap with known *cis*-regulatory elements such as riboswitches or RNA thermometers. Based on these criteria, we reassigned 20 noncoding RNAs to different sub-classes, while 16 candidates were removed completely (Supplementary Table 3). Updated annotations of the TSSs, terminators, operons, and noncoding RNAs can be accessed via Theta-Base 2.0 (www.helmholtz-hiri.de/en/datasets/bacteroides-v2).

### Prediction of invertible DNA regions

Invertible DNA regions were predicted using the PhaseFinder -locate pipeline as described in (Jiang et al., 2019): python PhaseFinder.py locate -f Bt_genome_plasmid.fa -t phasefinder.tab -g 15 85 -p. Briefly, inverted repeats were determined by allowing no mismatches for repeats of a maximum of 11 bp, one mismatch for repeats up to 13 bp, and two mismatches for repeats with lengths exceeding 19 bp. Homopolymeric inverted repeats were removed and the maximum GC content per inverted repeat was filtered to be between 15 and 85%.

### Differential gene expression analysis

Differential gene expression analysis was performed using the *R* package edgeR (3.38.2) (McCarthy et al., 2012; Robinson et al., 2010). Using filterByExpr function, the genes with a CPM of greater than 0.6635 (equivalent to ∼10 reads per sample) across all replicates in each growth condition (median library size of ∼15 million reads) were retained for differential analysis. While calling for contrasts (using makeContrast function), the analysis was sub-divided into two groups based on the respective control condition: all conditions with varying carbon sources including starvation were compared to glucose as a control, while all stress conditions were compared to the condition immediately prior to stress induction, i. e. a mid-exponential phase culture in TYG medium. Differential gene expression data across all conditions relative to their respective control condition can be found in Supplementary Table 7.

### Gene set annotation and enrichment analyses

We assembled a list of functionally annotated gene sets from the literature. We recovered annotation of PULs 01 to 96 from the PULdb (Terrapon et al., 2018); CPSs and CTns from (Xu et al., 2003); the genes transcribed from promoter motifs PM1 and PM2 from (Ryan et al., 2020a); regulons from RegPrecise v3.2 (Novichkov et al., 2013); the BT_4338-regulon from (Townsend et al., 2020); annotated KEGG pathways and modules from the KEGG database (accessed on 01/Dec/22); GO terms from Uniprot (accessed on 25/Nov/21); predicted KEGG modules and pathways and GO terms from an eggNOG v5.0 (Huerta-Cepas et al., 2019) annotation of the *B. thetaiotaomicron* genome. Gene set enrichment analysis was performed with the fgsea *R* package over all gene sets with >9 genes, except for PULs, which were retained irrespective of their gene number. Genes were ranked based on the -log10(PValue)*sign(fold change) metric.

### Northern blotting

Northern blotting was performed as described previously (Ryan et al., 2020a). In short, total RNA (2.5-10 µg) was electrophoretically resolved on a 6% (vol vol^-1^) polyacrylamide (PAA) gel containing 7 M urea and electro-blotted onto a membrane (Hybond XL, Amersham) at 50 V, 4 °C for 1 h. The blots were then probed with gene-specific ^32^P-labeled oligonucleotides in Hybri-Quick buffer (Carl Roth AG) at 42°C and subsequently exposed to a phosphor screen as required. Images were visualized using a phosphorimager (FLA-3000 Series, Fuji).

### Reanalysis of transposon insertion sequencing data

We reanalyzed fitness data from a comprehensive *B. thetaiotaomicron* transposon mutant library that probed a suite of over 490 different conditions including 48 different carbon sources and 56 stress-inducing compounds, in the context of our extensive ncRNA annotation (Liu et al., 2021). This was done with the primary objective of identifying and possibly correlating gene expression from our transcriptomic dataset with mutant fitness data and thereby allow us to draw biologically meaningful conclusions. In order to further streamline our analysis, we focused exclusively on independently encoded intergenic sRNAs since phenotypes pertaining to such mutants would likely not involve polar effects. Consequently, of the 135 intergenic sRNAs identified in *B. thetaiotaomicron* till date, we obtained phenotypes for 95 sRNAs (∼70%), of which 32 were associated with statistically significant fitness (|t|>4) in at least one “successful” experiment (Supplementary Table 5). The fitness of a gene is defined as the average log2 change in relative abundance of its mutants (|fit|). Negative and positive values mean that the sRNA mutants were less or more fit, respectively, than the average strain. A “successful” experiment requires that each gene is represented by a sufficient number of barcode counts (Wetmore et al., 2015). Experiments with ‘jackpot’ effects whereby the disruption of an sRNA results in a large competitive advantage versus the other mutants in the pool, were retained, but specifically labeled as “strong” phenotypes (|fit| > 2 and |t| > 5) (Supplementary Table 5). A third category namely “combined” comprised those phenotypes that were both “strong” and “significant”, as per the above criteria.

### Launch of Theta-Base 2.0

Theta-Base 2.0 (www.helmholtz-hiri.de/en/datasets/bacteroides-v2) was created with Flask (Grinberg, 2018) (back-end) and Vue.js (https://vuejs.org/guide/introduction.html) (front-end), storing underlying visualization and expression data using MongoDB. The Clustergrammer plugin utilizes the API from the Ma’ayan lab (Fernandez et al., 2017), while the heat map plugin follows the same front-end and back-end architecture as the main site. Gene set annotations (GO Terms, KEGG pathways and modules, polysaccharide utilization loci, capsular polysaccharides, conjugative transposons, promoter motifs, and known regulons) were prepared as described in the section “Gene set annotation and enrichment analyses” and can be found in Supplementary Table 8. The sRNA fitness dataset was adapted from Supplementary Table 5. Deployment of the back-end and front-end use Gunicorn (https://readthedocs.org/projects/gunicorn-docs) and Nginx (Reese, 2008).

*B. thetaiotaomicron* datasets can be manually selected by first clicking the “Bacteroides Theta” tab followed by providing a “Title” and then selecting an appropriate dataset using the dropdown menu. Users can select from a choice of expression data normalized in CPM, Log_2_FC (compared to control conditions), the entire dataset, or sRNA fitness data. As an option, columns of interest can be further customized using the “Select columns” box and subsequently clicking the “Add” tab. Users also have the option to add their own data as outlined in additional tabs, such as uploading a delimited file.

The resulting data tables are displayed in the browser and can be filtered or transformed. For example, the “Filter” button allows data tables to be filtered using keywords with prompts to make the search process seamless. The “Functional annotation” button permits the user to select from a large number of preset manually curated gene lists such as “Go Term”, “KEGG pathway”, “PUL”, “CPS”, and “CTn”, to name a few. A third button, “ncRNA” permits selection of manually curated noncoding RNA gene sets for example, “High confidence intergenic sRNAs”, “Intergenic sRNAs”, and “*Cis*-antisense RNAs”. Once the desired genes have been filtered, they may be transformed by clicking the “Transform Data” button such as rounding values, log conversion, calculating transcript per million (TPM), and others. Once the appropriate datasets have been defined and loaded by the user, they can be further examined using three modes, namely “Heatmap”, “Clustergrammer”, and “JBrowse”. 2D and 3D heat maps can be generated using the “Heatmap” function. Note that the heat map defaults to a 3D option, but users can manually switch to the 2D option. Heat maps can be customized by using the menu on the left that permits changes to the color gradients and overall structure. Customized heat maps can be downloaded in SVG or PNG formats using the download tab. Alternatively, if users wish to probe the selected datasets and investigate relationships by clustering according to genes or conditions, “Clustergrammar” is recommended. Selecting this tab generates a 2D dynamic heat map of the data that can be further investigated using the menu on the left. Currently, the tool permits a maximum of 200 rows to be loaded and users will be prompted if more rows are selected. Customized heat maps and data tables can be downloaded using the “Take snapshot” and “Download matrix” buttons, respectively. For a detailed view of normalized coverage plots for the investigated conditions, in addition to those published in the first iteration of Theta-Base (Ryan et al., 2020a), users can select the JBrowse button. Users are free to select from a range of updated annotations displaying high-resolution maps for non-coding RNAs (ncRNA), transcription start sites (TSSv3), terminators (term_v2), operons (Operon_structure), a transposon insertion map related to the fitness data (Tn_insertions), and invertons (Inverted_repeats).

On the top right of the website there are four buttons. The “padlock button” locks the current state of the site, allowing users to copy their URL and share with colleagues. The next button allows users to download the currently selected dataset as an Excel or a delimited file (such as .csv). The new document button will re-load the website so users can select a new dataset. The help button when clicked will provide pop-over text explaining various features of the site.

### Correlation analysis of sRNA expression and fitness of the corresponding transposon mutants

Among the 15 experimental conditions for RNA-seq in this study, five metabolic (namely D-glucose, L-arabinose, D-xylose, D-maltose monohydrate, GlcNAc) and four stress-related conditions (deoxycholate, bile salts, pH 6, aerobic shaking) were similar to conditions previously covered in the comprehensive TIS data (Liu et al., 2021). To assess whether sRNA expression showed any relationship with the fitness scores of the corresponding sRNA disruption mutants, we calculated relative expression (read count in terms of z-score) for each sRNA using the mean read counts per million (CPM) and standard deviation of the CPM across all CPMs in the 15 growth conditions. The plot in Supplementary Fig. S7b shows the z-scores of the 95 selected sRNAs plotted against their respective TIS fitness scores for the above nine growth conditions.

### Antibiotics growth curve analyses and agar strip assays

Bacterial growth curves were performed by inoculating each a single colony of AWS-003 (WT) and AWS-0029 (Δ*masB*) into 5 mL of TYG medium and incubation overnight under anaerobic conditions. These cultures were then sub-cultured (1:100 dilution) in 2 mL of TYG medium containing doxycycline (0.016 µg mL^-1^, Sigma-Aldrich), tetracycline (0.05 µg mL^-1^, AppliChem), or a water control. The samples (200 µL) were incubated in a 96-well flat-bottom plate (Nunclon) at 37°C with continuous shaking (double orbital) in a microplate spectrophotometer (BioTek Epoch 2). Optical densities were recorded every 20 min. The assay was performed in each three biological replicates.

Antibiotics strip assays were performed by dipping a sterile cotton swab into TYG-overnight cultures of AWS-003 or AWS-0029 and streaking on BHIS agar plates containing strips for doxycycline (HIMEDIA, EM103) and tetracycline (HIMEDIA, EM056). The plates were incubated anaerobically for 48 h at 37°C and images taken. The minimal inhibitory concentrations (MICs) were derived from the positions where the inhibition ellipses intersected the strips.

### Gene coexpression analyses

Correlation of expression of all *B. thetaiotaomicron* genes across all the profiled carbon source and stress conditions was calculated by generating a correlation matrix (Pearson’s correlation score) of the z-scores of the CPM values of each gene. To identify the correlation in expression between our gene sets (see “Gene set annotation and enrichment analyses”) and a given gene of interest (MasB in Fig. 5a and *BT_1675* in Supplementary Fig. S10), the median of the correlation values between all genes within a gene set and the gene of interest was calculated. Gene sets composed of <10 operons were excluded from this analysis.

### MS2 affinity purification and sequencing

A *B. thetaiotaomicron* Δ*masB* mutant complemented with either MS2-MasB (AWS-062) or untagged MasB (AWS-036) was diluted 1:100 in 50 mL of TYG medium from an overnight culture grown anaerobically at 37°C. At OD_600_ of 2.0, expression of MS2-tagged MasB and untagged MasB was induced by the addition of 200 ng/mL of anhydrotetracycline (aTC). After two additional hours of growth at 37°C, 25 mL of the cultures (equivalent to 90 ODs) were collected, centrifuged for 20 min at 4,500 rpm at 4°C, and snap-frozen in liquid nitrogen. MS2 pulldown and RNA purification was performed as described in (Correia Santos et al., 2021), yet with slight modifications to adapt the protocol to *Bacteroides*. Specifically, the column was washed only six (instead of eight) times with buffer A prior to elution. Elution itself was then induced with 600 (rather than 300) µL of elution buffer.

For library preparation (at Vertis Biotechnologie AG), the RNA samples were first fragmented using ultrasound (4 pulses of each 30 s at 4°C). Then, an oligonucleotide adapter was ligated to the 3’ ends of the RNA molecules. First-strand cDNA synthesis was performed using M-MLV reverse transcriptase and the 3’ adapter as primer. The first-strand cDNA was purified and the 5’ Illumina TruSeq sequencing adapter was ligated to the 3’ end of the antisense cDNA. The resulting cDNA was PCR-amplified to about 10-20 ng/μL using a high fidelity DNA polymerase. The cDNA was purified using the Agencourt AMPure XP kit (Beckman Coulter Genomics) and was analyzed by capillary electrophoresis. For Illumina sequencing, the samples were pooled in approximately equimolar amounts. In order to deplete cDNAs derived from 5S rRNA, the cDNA pool was digested using probes specific for bacterial 5S sequences and Cas9 endonuclease. Afterwards, the cDNA pool was fractionated in the size range of 200–600 bp using a preparative agarose gel. An aliquot of the size-fractionated pool was analyzed by capillary electrophoresis. The cDNA pool was sequenced on an Illumina NextSeq 500 system using 2×150 bp read-length.

Generated reads were quality-checked using FastQC (v0.11.8) and adapters were trimmed using bbduk with the following parameters: qtrim=r trimq=10 ktrim=r ref=bbmap/ressources/adapters.fa k=23 mink=11 hdist=1 tpe tbo. BBmap was used to map reads to the *B. thetaiotaomicron* VPI-5482 reference genome (NC_004663.1) and plasmid (NC_004703.1) as well as to the MS2-MasB sequence. Read quantification was performed using featureCounts (2.0.1). Differential gene expression analysis between the MS2-MasB and untagged samples was conducted using edgeR (Chen et al., 2016) in combination with RUVSeq (Risso et al., 2014) to estimate the factor of unwanted variation using replicate sample (RUVs) with correction factor k=1.

### IntaRNA prediction of sRNA–mRNA interactions

*In-silico* interaction prediction between MasB and its putative mRNA targets, *fusA2* and *BT_1675*, was performed with the help of IntaRNA (Mann et al., 2017) using the Vienna RNA package (2.4.14 and boost 1.7) at default settings along with the output flag (--out=pMinE:FILE.csv) to generate minimal energy values for intermolecular index pairs. For visualization, the resulting values were plotted in form of a heat map in *R* (v4.2).

### In*-*vitro transcription and radiolabeling of RNA

DNA templates for *in-vitro* transcription were amplified using genomic DNA and primer pairs carrying a T7 promoter (Supplementary Table 1). The *in-vitro* transcription reaction was performed using the MEGAscript T7 kit (ThermoFisher Scientific) followed by DNase I digestion (1 U, 37°C for 15 min). RNA products were then excised from a 6% (vol vol−1) PAA-7M urea gel by comparison to a LowRange RNA ladder (ThermoFisher Scientific) and eluted overnight in elution buffer (0.1 M NaOAc, 0.1% SDS, 10 mM EDTA) on a thermoblock at 8°C and 1,400 rpm. The next day, the RNA was precipitated in an ethanol:NaOAc (30:1) mix, washed with 75% ethanol, and resuspended in 20 µL water (65°C for 5 min).

Radioactive labelling of the *in-vitro*-transcribed RNA was carried out by first dephosphorylating 50 pmol of RNA with 25 U of calf intestine alkaline phosphatase (NEB) in a 50 µL reaction and incubated at 37°C for 1 h. The dephosphorylated RNA was extracted using phenol:cholorform:isoamylalcohol (P:C:I, 25:24:1) and precipitated as described above. 20 pmol of this RNA was 5’-end-labelled (20 µCi of ^32^P-γATP) using 1 U of polynucleotide kinase (NEB) at 37°C for 1 h in a 20 µL reaction. The labelled RNA was then purified using a G-50 column (GE Healthcare) and extracted from a PAA gel as described above.

### Electrophoretic mobility shift assay

EMSA was carried out in a reaction volume of 10 μL, containing 1x RNA structure buffer (SB; Ambion), 1 μg yeast RNA (∼4 μM final concentration), 5′-end-labeled MasB RNA (4 nM final concentration) and an mRNA segment of 137 nt in length, spanning the predicted MasB-target site within *BT_1675* (see Supplementary Fig. S9b) at final concentrations: 0; 8; 16; 32; 64; 128; 256; 512; and 1,024 nM. Following incubation at 37 °C for 1 h, 3 μL of 5x native loading dye (0.2% bromophenol blue, 0.5x TBE, 50% glycerol) were added to each tube. All samples were loaded on a native 6% (vol vol−1) PAA gel in 0.5x TBE buffer at 4°C and run at 300 V for 3 h at this temperature. The gel was then dried, exposed, and visualized using a phosphorimager (FLA-3000 Series, Fuji). The experiment was repeated three times and quantified using ImageJ v1.52s (Schneider et al., 2012) and GraphPad Prism version 9 for Windows, GraphPad Software, San Diego, California USA, www.graphpad.com. The dissociation constant (K*_d_*) was calculated via the one site -- specific binding formula:

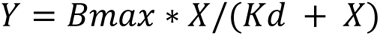

where *Y* : specific binding; *X* : concentration of radio-ligand; *B_max_*: maximum binding in the same unit as *Y*; *Kd*: in the same unit as *X*.

### Quantitative real-time PCR

For qRT-PCR assays, Δ*tdk B. thetaiotaomicron* (AWS-003; referred to as wild-type in Fig. 5), Δ*masB* (AWS-029), and *masB*^+^ (AWS-036) were grown anaerobically overnight at 37°C in 5 mL of TYG medium, then sub-cultured 1:100 in 25 mL of TYG, and induced with 200 ng/mL of aTC. ∼4 OD equivalents of samples were collected at early exponential phase (OD_600_ = 0.3), mid exponential phase (OD_600_ = 2.0), and late exponential phase (OD_600_ = 3.7) for RNA extraction as described above. For the starvation condition, the same strains were grown anaerobically for 24 h at 37°C in 5 mL of minimal medium supplemented with 0.5% glucose, then sub-cultured 1:100 in 25 mL of 0.5% glucose-containing minimal medium supplemented with 200 ng/mL aTC. At OD_600_ = 2.0, the cultures were centrifuged, the supernatant was discarded, and the pellet resuspended in minimal media without carbon source, and incubated anaerobically at 37°C for another two hours before ∼4 ODs of the samples were collected for RNA extraction. qRT-PCR reactions were performed as described in (Ryan et al., 2020a). Four technical replicates per each biological replicate were pipetted and plates analyzed on a QuantStudio5 instrument (ThermoFisher).

---

**Supplementary Figure S1:**
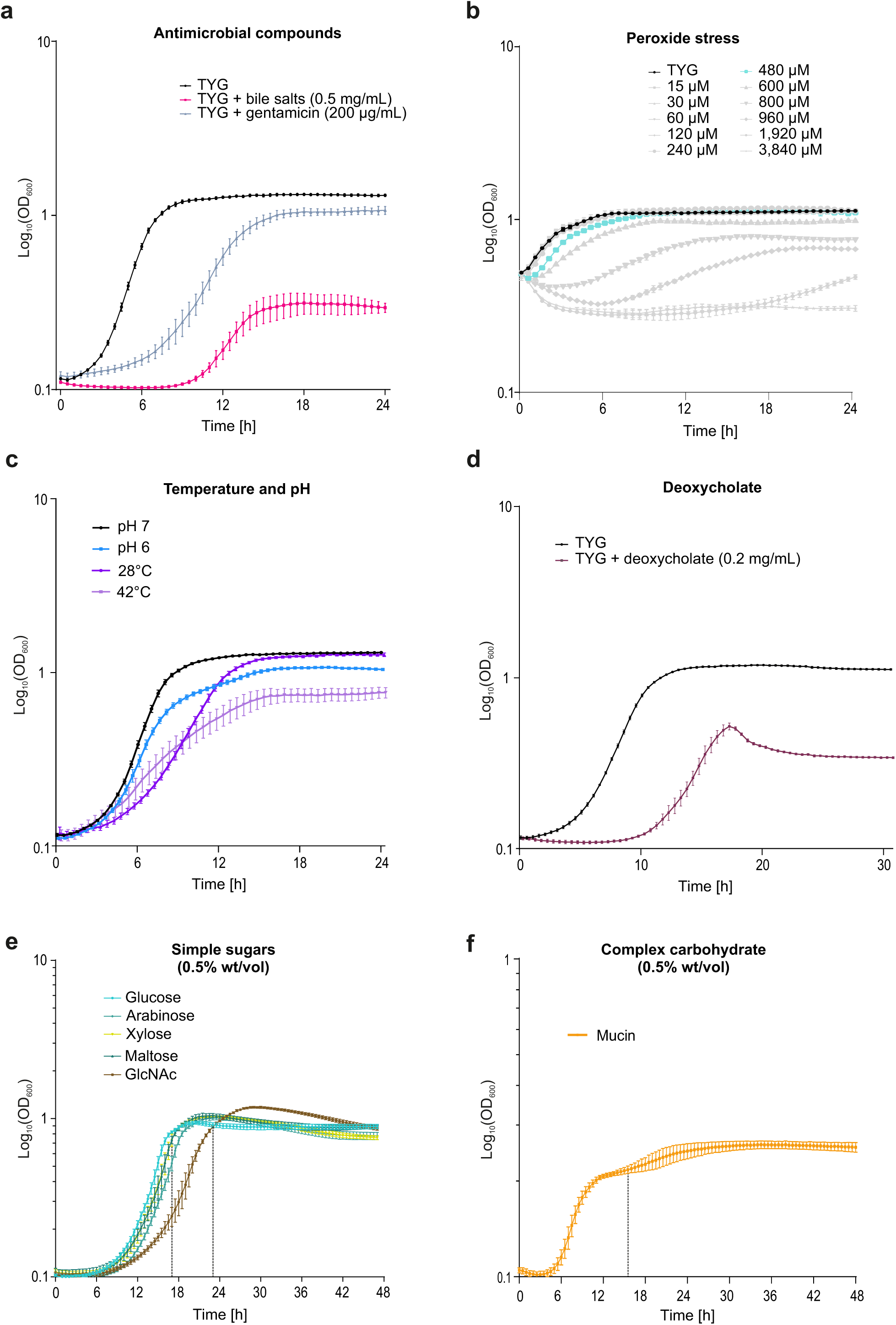
*B. thetaiotaomicron* growth curves over the profiled *in vitro* conditions. **a–d,** Growth in rich TYG medium upon constant exposure to the indicated environmental stresses. **e, f,** Growth in minimal medium with the indicated carbohydrates— simple sugars (**e**) and porcine mucin (**f**)—added as sole carbon sources. In each case, growth curves and error bars denote the mean +/- SD from each three biological replicates. The dashed vertical lines in panels **e** and **f** denote the time points of sampling.

**Supplementary Figure S2:**
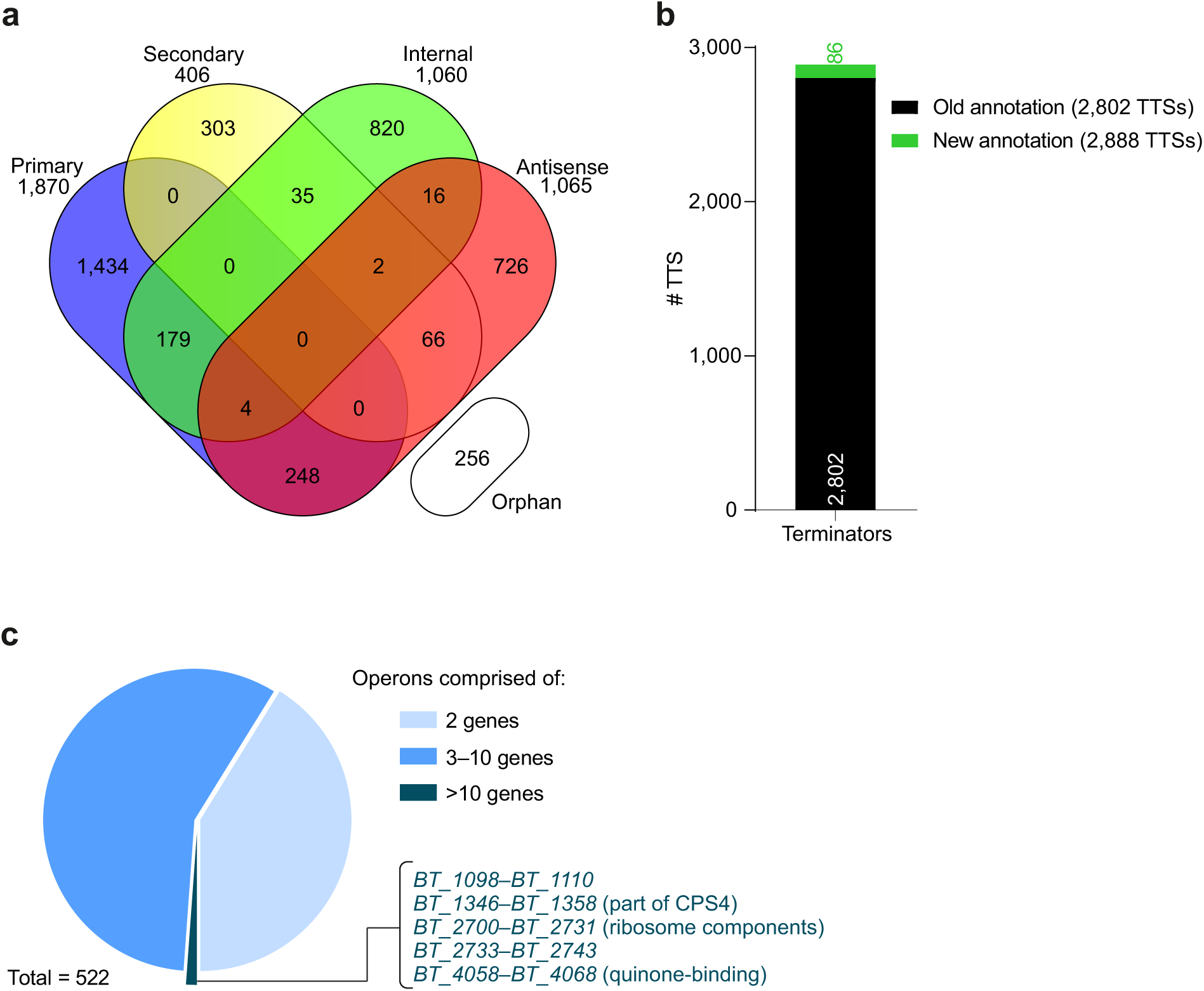
Updated transcription start site annotations, intrinsic terminator annotations and operon structures of *B. thetaiotaomicron*. **a,** Venn diagram showing the updated numbers of transcription start site categories. **b,** Refined TTS annotations. **c,** Operon structure prediction. Operons encompassing more than 10 genes are labeled by name.

**Supplementary Figure S3:**
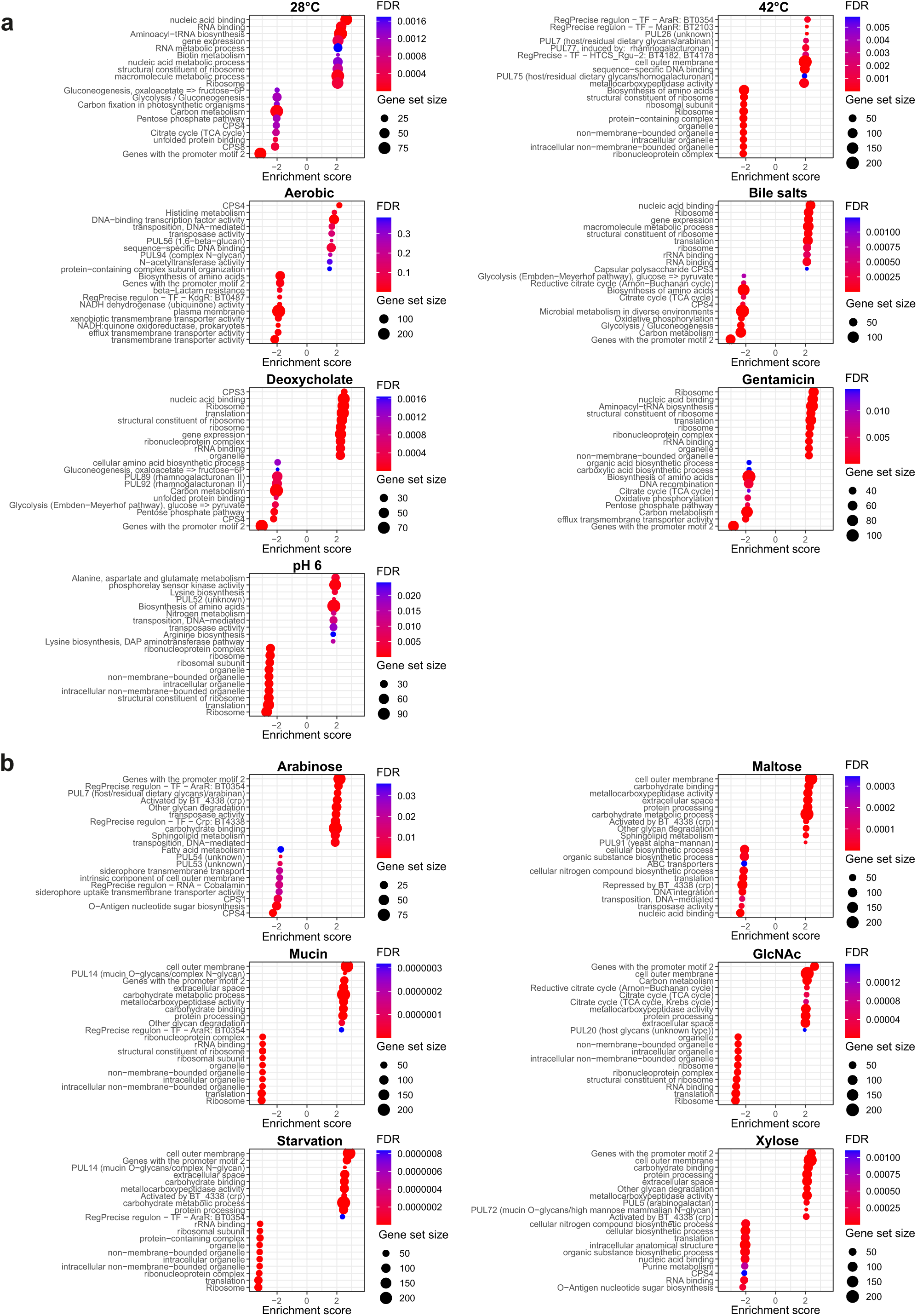
Gene set enrichment analysis. Top 10 enriched (normalized enrichment score >0) and depleted (normalized enrichment score <0) gene sets in the stress (**a**) and carbon source conditions (**b**). Gene sets derive from the custom gene set list described in the Methods section under “Gene set annotation and enrichment analyses”. Each gene set is represented by a circle whose size is proportional to the number of included genes (gene set size). Known inducers of PULs are included in brackets after the PUL name.

**Supplementary Figure S4:**
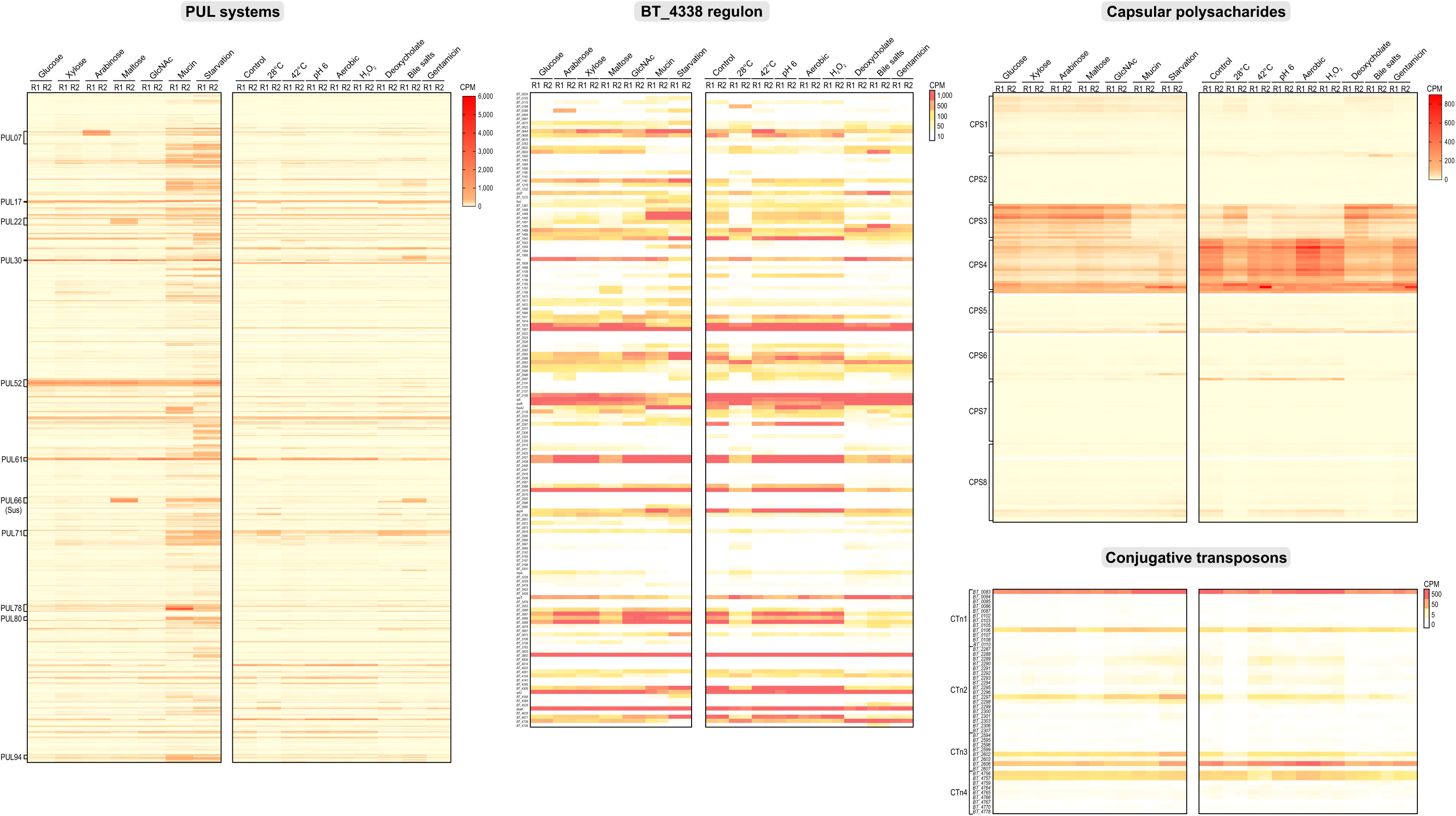
Transcript levels of all annotated *B. thetaiotaomicron* PUL systems, BT4338 regulon members, capsular polysaccharides, and conjugative transposons. For each condition and each replicate, the read counts per million (CPM) mapped to individual PUL genes (as inferred from PULDB (Terrapon et al., 2018)), to individual members of the BT4338 regulon as inferred from (Supplementary Table S3B in (Townsend et al., 2020)), to CPS genes, and to CTn loci are plotted. Selected, strongly affected PULs and CPSs are labeled and re-plotted in Fig. 2c.

**Supplementary Figure S5:**
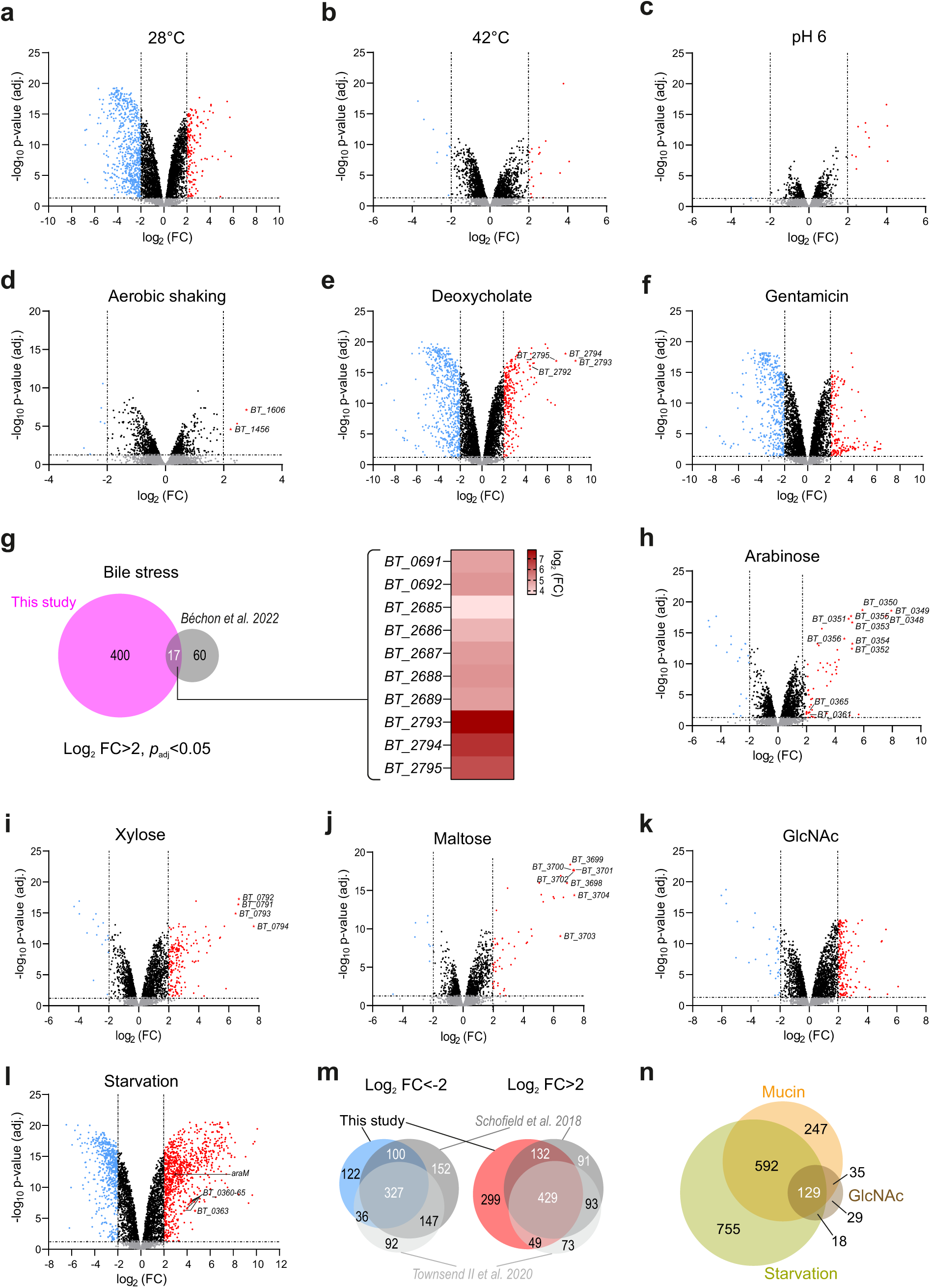
Pair-wise comparison of stress- and carbon source-specific gene expression changes. **a–g,** Volcano plots report differential expression of *B. thetaiotaomicron* exposed to the indicated stresses (x-axis) over significance (y-axis). **a,** Cold vs. TYG control. **b,** Heat vs. TYG control. **c,** Acidic vs. TYG control. **d,** Aerobic shaking vs. TYG control. The genes for cytochrome C peroxidase (*BT_1606*) and thioredoxin (*BT_1456*) are labeled. **e,** Deoxycholate vs. TYG control. Genes belonging to the *BT_2792-BT_2795* operon (https://doi.org/10.1101/573055) are labeled. **f,** Gentamicin vs. TYG control. **g,** Venn diagrams display the overlap of bile salts-specific gene expression (significantly upregulated) between our dataset and that obtained in (Bechon et al., 2022). The associated heat map depicts highly upregulated operons including components of two efflux systems (*BT_2793-BT_2795*, *BT_2685-BT_2689*) as well as an outer membrane protein and a calcineurin superfamily phosphohydrolase (*BT_0691-BT_0692*). **h–n,** Volcano plots report differential expression of *B. thetaiotaomicron* feeding on the indicated carbon sources (x-axis) over significance (y-axis). **h,** Arabinose vs. glucose. Genes belonging to the arabinose utilization operon *BT_0348-BT_0369* (Schwalm et al., 2016) are indicated. **i,** Xylose vs. glucose. Genes belonging to the xylose utilization operon *BT_0791-BT_0794* (Townsend et al., 2020) are indicated. **j,** Maltose vs. glucose. Genes belonging to the starch utilization system (*sus*) operon *BT_3704-BT_3698* (Cho et al., 2001; Reeves et al., 1997) are indicated. **k,** *N*-acetyl-D-glucosamine vs. glucose. **l,** Starvation vs. TYG. **m,** Venn diagram denotes the overlap of starvation-induced up- or downregulations observed here and in previous studies (Schofield et al., 2018; Townsend et al., 2020). **n,** Venn diagram denotes the overlap in the regulated gene sets between *N*-acetyl-D-glucosamine or mucin-consuming bacteria, and starved cultures.

**Supplementary Figure S6:**
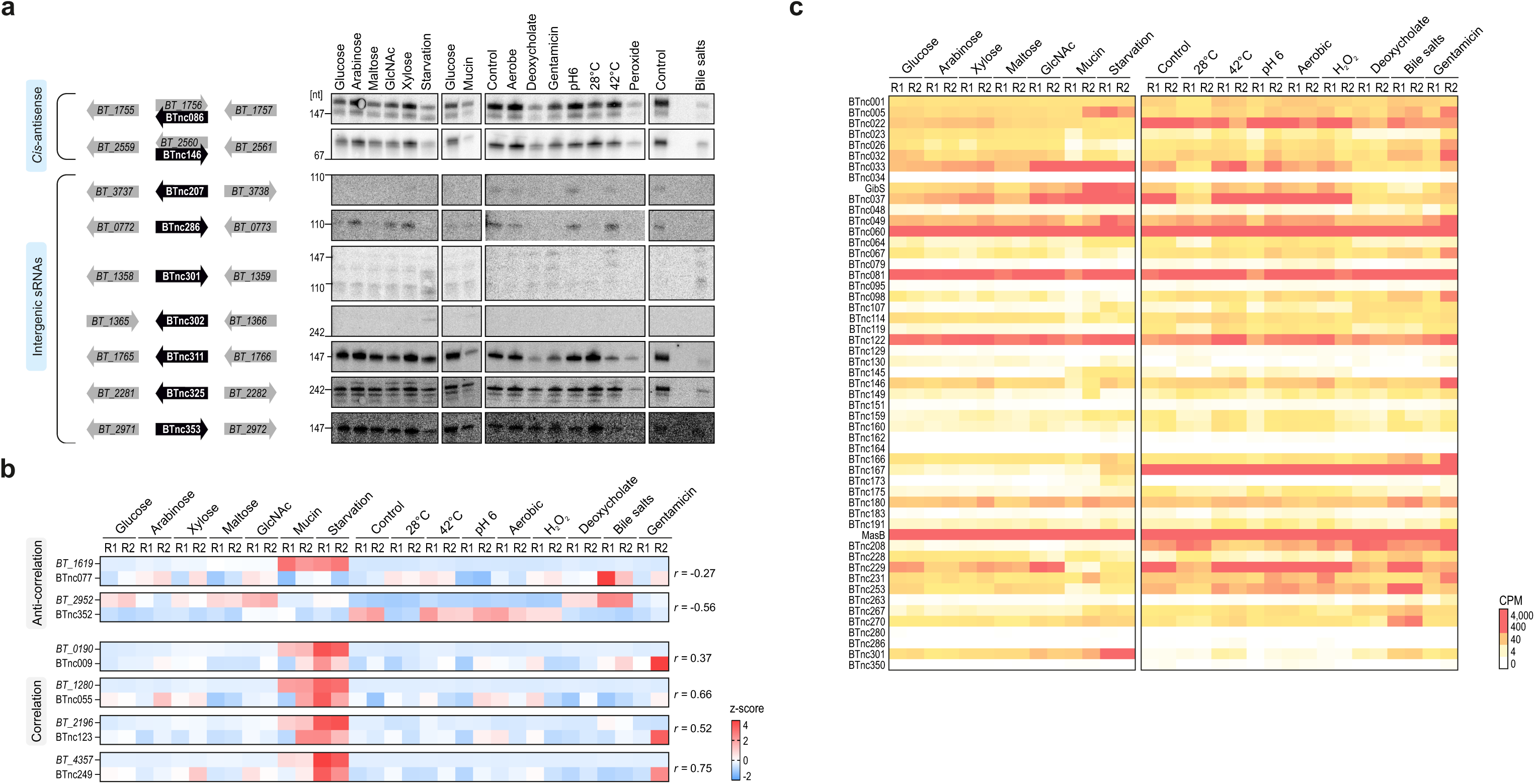
Expression of *B. thetaiotaomicron cis*- and *trans*-encoded noncoding RNAs. **a,** Northern blot-based validation of newly predicted noncoding RNA candidates. **b,** Log_2_FC values of PUL-associated antisense RNAs and their respectively overlapping PUL gene (*susC* homologue) reveal patterns of either correlation or anti-correlation. **c,** Overall abundance (CPM, counts per million) of ‘canonical’ sRNAs, defined as intergenic sRNAs harboring both an experimentally identified TSS and a predicted intrinsic terminator, across the suite of experimental conditions and replicates.

**Supplementary Figure S7:**
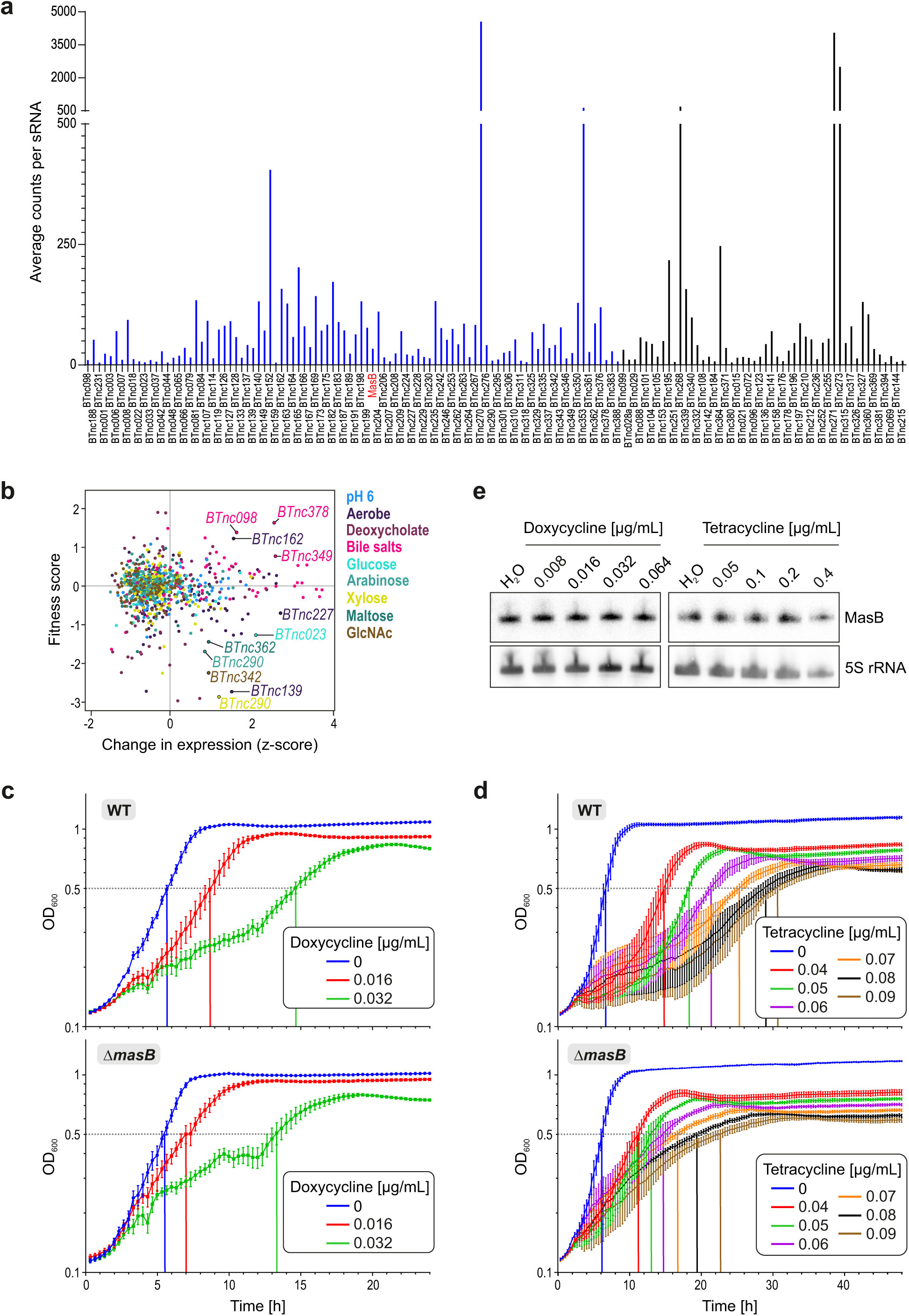
TIS screen for phenotypes associated with *B. thetaiotaomicron* sRNAs. **a,** Average number of transposon insertions per ncRNA gene in the mutant library across the range of different sample conditions. Intergenic *B. thetaiotaomicron* sRNAs are shown in blue, and 5’- or 3’-derived, or intra-operonic sRNA candidates in black. In this study, we focused on only the intergenic sRNA mutants. **b,** Associating sRNA expression profiles with their contribution to bacterial fitness. For the nine experimental conditions shared between transcriptome profiling and the TIS screen, relative expression (z-scores) of all 81 intergenic sRNAs hit by a transposon insertion is plotted over the change in fitness of the corresponding mutants. **c, d,** Growth curves of *B. thetaiotaomicron* isogenic wild-type (upper) and Δ*masB* (lower) in TYG supplemented with increasing concentrations of doxycycline (**c**) or tetracycline (**d**). Plotted are the means +/-SD from each three biological replicate experiments, that each comprised technical duplicates. Indicated with dotted lines are the times to reach an OD_600_ of 0.5 for each strain and treatment. **e,** Antibiotics exposure does not majorly influence MasB steady-state levels. Northern blot on total RNA samples derived from *B. thetaiotaomicron* cultures exposed for 2 h to increasing concentrations of either doxycycline or tetracycline, relative to RNA from vehicle (water)-treated control cultures. 5S rRNA was the loading control.

**Supplementary Figure S8:**
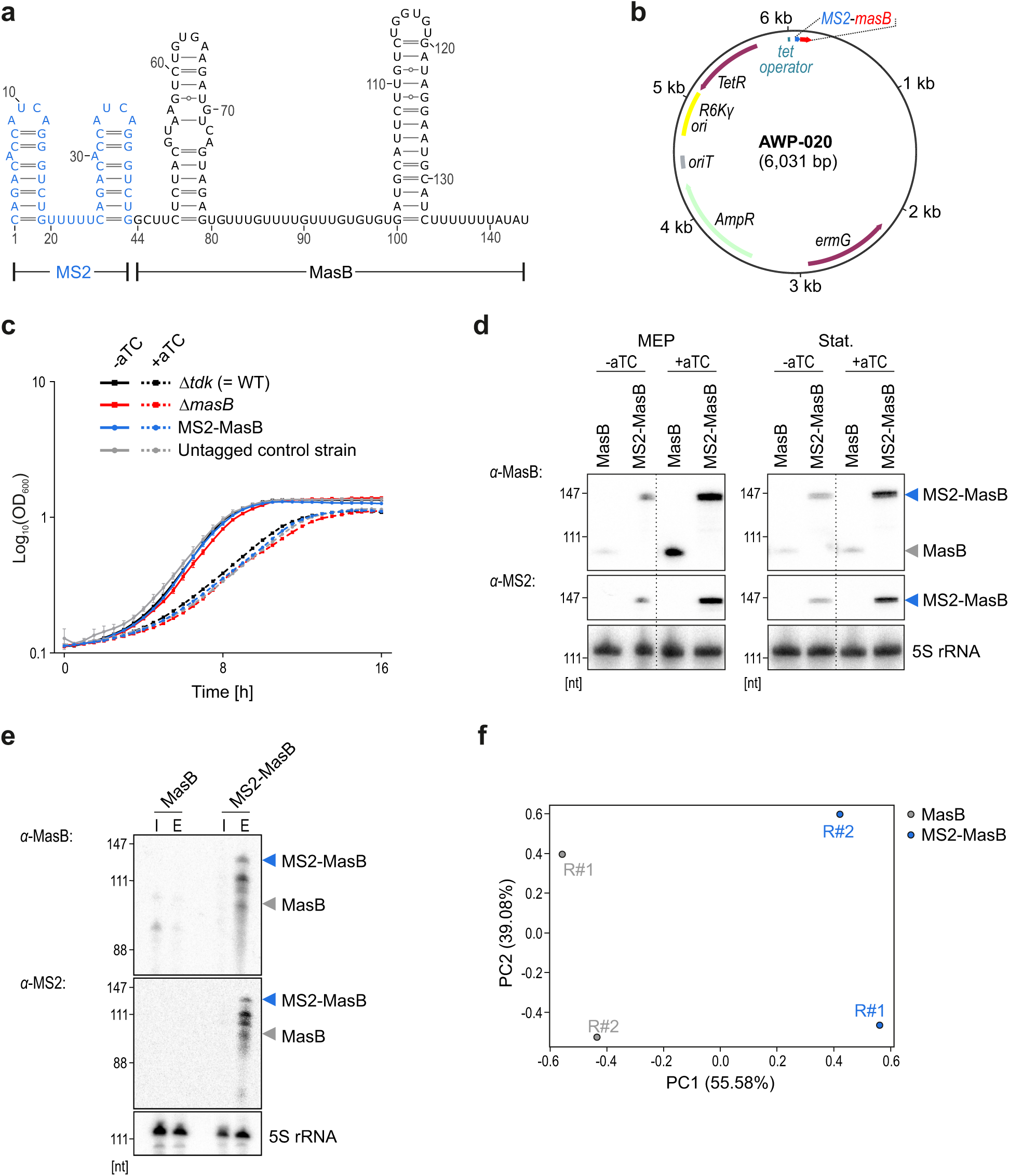
Establishment of MAPS for *B. thetaiotaomicron* sRNA MasB. *In-silico* prediction of the secondary structure of a fusion of the 5’ end of MasB to the MS2 aptamer using the RNAfold WebServer (Lorenz et al., 2011). **b,** Plasmid map of AWP-020 containing the anhydrotetracycline-inducible MS2-MasB construct. **c,** Growth curves of the indicated strains in TYG medium in the absence (solid line) or presence (dotted line) of anhydrotetracycline (aTC; 200 ng/mL) as an inducer of MasB expression. Plotted values are the means of three biological replicates with error bars indicating the standard deviation. **d,** Northern blot to probe MS2-MasB in the strains used for MAPS grown in TYG to mid-exponential phase (MEP, ∼7 h) or stationary phase (∼10 h) in the absence or presence of anhydrotetracycline induction (for 2 h). **e,** Northern blot of the input (I) and eluate (E) fractions of the MAPS experiment, probed for MasB or MS2. 5S rRNA served as the loading control. An enrichment of MS2-MasB in the eluate compared to the input and to the samples derived from the untagged control strain demonstrates efficient capture and pull-down of MasB. **f,** Principal component analysis plot of the sequencing data revealed segregation between the two biological replicates derived from the pull-down of MS2-MasB (blue) and those from the control strain (grey).

**Supplementary Figure S9:**
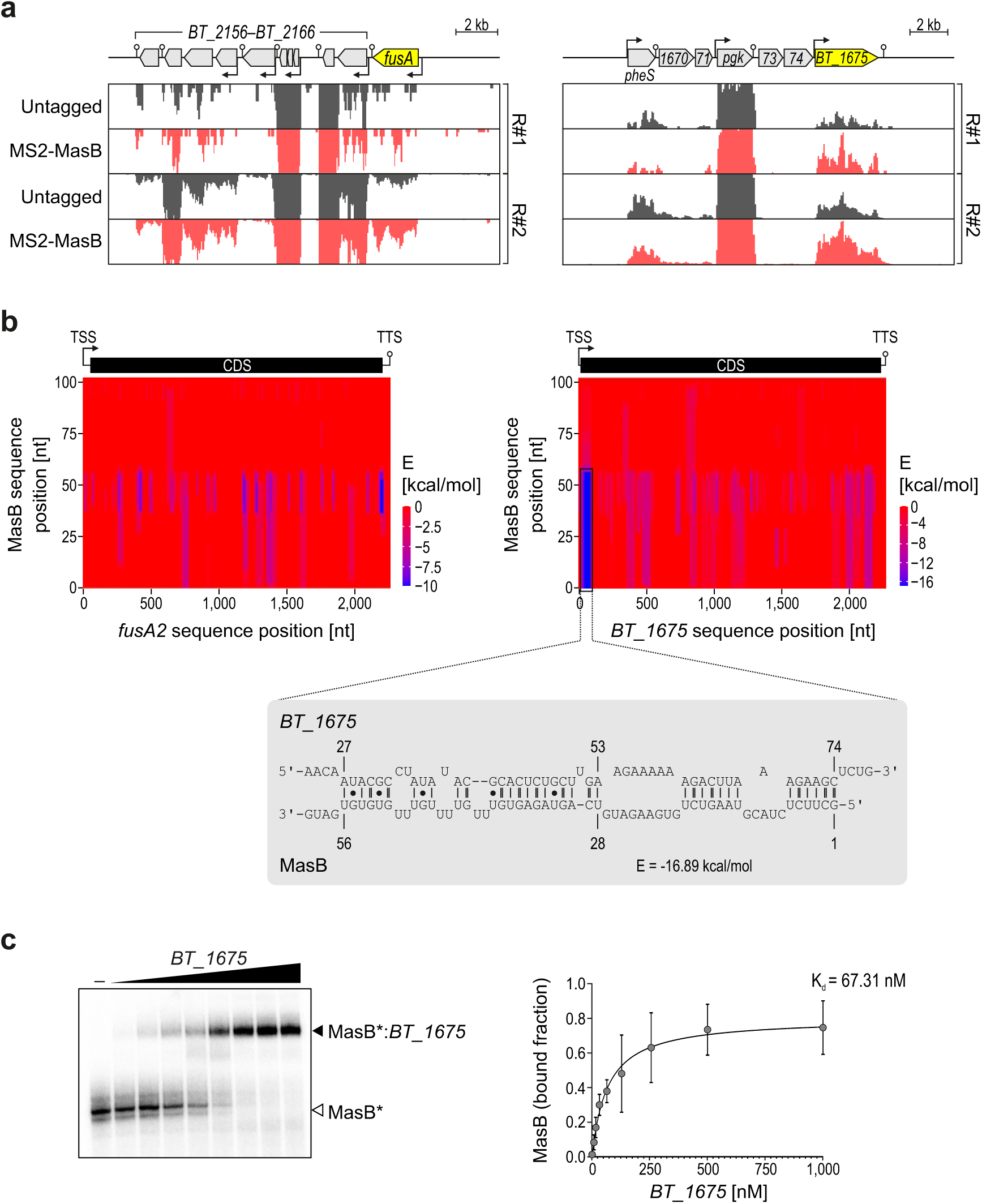
Validation of the direct interaction between MasB and *BT_1675* mRNA. **a,** MAPS coverage plots across the *fusA2* and *BT_1675* loci. R#1 and #2 are biological replicates. **b,** *In-silico* prediction of putative MasB interaction sites within MAPS-derived target candidates. Upper: the heat maps display the position-wise minimal energy (E) profiles of *fusA2* or *BT_1675* mRNAs (x-axes), respectively, with MasB (y-axis) as retrieved from IntaRNA (Mann et al., 2017). Full-length sequences from TSS to TTS have been queried and the positions are relative to the TSS in each case. Lower: depiction of the top-ranked interaction region within *BT_1675* mRNA at the nucleotide-level. Here, numbers refer to nucleotide positions relative to the translational start codon (in case of the mRNA) or the TSS (for MasB). **c,** EMSAs to validate the predicted MasB binding within *BT_1675. In-vitro*-transcribed and radio-labeled MasB RNA was incubated with increasing concentrations of a 137 nt-long 5′ segment of *BT_1675* mRNA. A representative blot is shown alongside the dissociation curve for three technical replicates. Arrowheads in white or black indicate free or bound MasB, respectively.

**Supplementary Figure S10:**
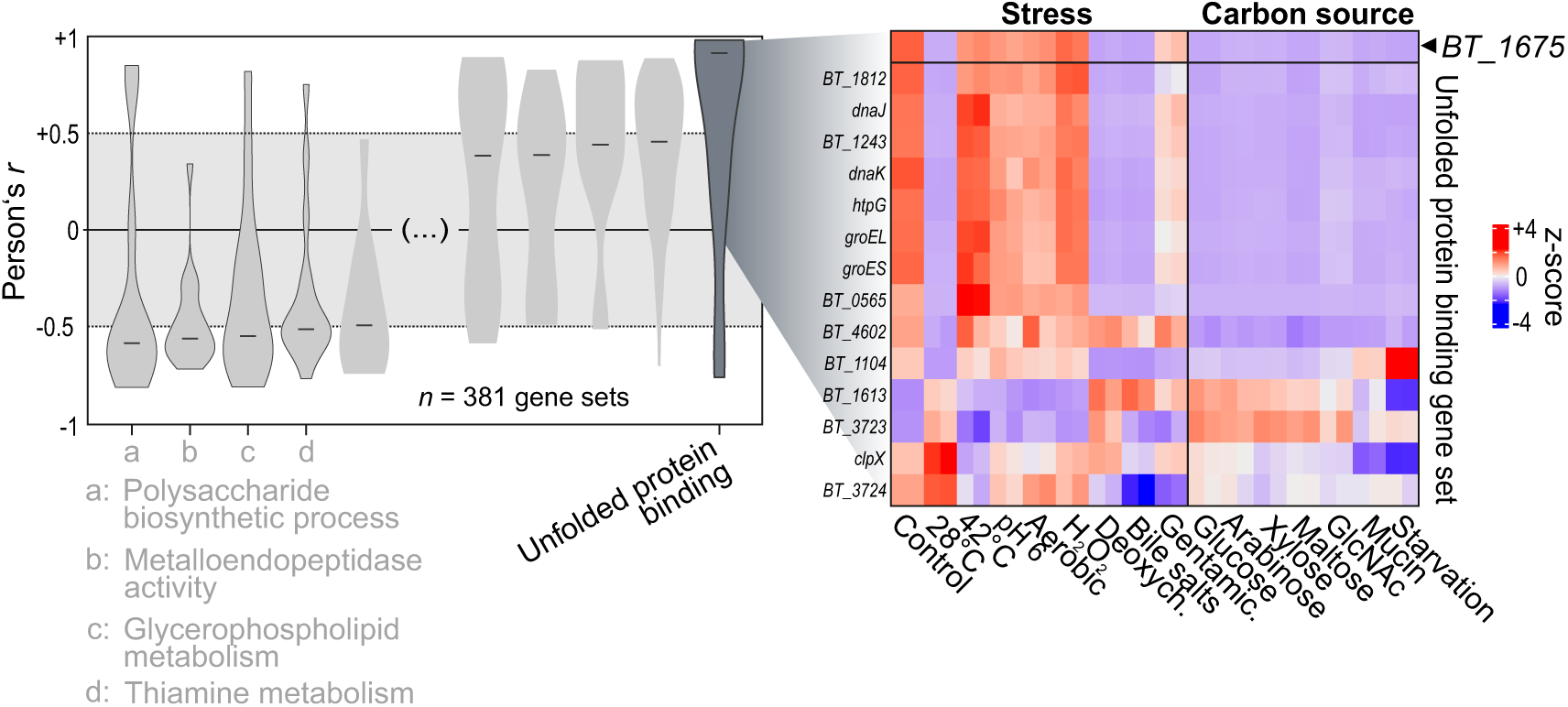
Co-expression analysis of *BT_1675*. **a** Left: violin plots of Pearson’s correlation coefficients between MasB expression and that of annotated *Bacteroides* gene sets (≥10 individual transcriptional units) sorted from left (lowest *r*) to right (highest *r*). Out of all 381 gene sets included in this analysis, the 5 top positively and negatively correlated (highest and lowest median r, respectively) are shown. The GO term “Unfolded protein binding” ranked first in absolute correlation. Gene sets with |median *r*| >0.5 are named below the plot. Right: heat map showing the expression of *BT_1675* and “Unfolded protein binding” genes across the set of 15 different experimental conditions.

## ACKNOWLEDGEMENT

We thank Sarah Reichardt for technical support, Ana-Rita Brochado (University of Würzburg) for kindly sharing antibiotics, Jörg Vogel and Anke Sparmann for constructive feedback on this manuscript, and all members of the Westermann lab for fruitful discussions. We would also like to thank Michael Kütt for his help when setting up the Theta-Base 2.0 website, Laura Jenniches for help with JBrowse, Sara Correia Santos for helpful discussions about MAPS,, and Morgan Price (LBNL) for calculating gene fitness scores. E.B. and T.F.C. were recipients of fellowships from the HIRI graduate training program “RNA & Infection”. A.M.D acknowledges support from the U.S. National Institutes of Health grant RM1 GM135102. Research in the Westermann laboratory is supported by the German Research Foundation (DFG; Individual Research Grant We6689/1-1) and the European Research Council (ERC Starting Grant “GUT-CHECK”).

## Notes

http://www.helmholtz-hiri.de/en/datasets/bacteroides-v2

### Competing Interest Statement

The authors have declared no competing interest.

